# Resolving chaperone-assisted protein folding on the ribosome at the peptide level

**DOI:** 10.1101/2022.09.23.509153

**Authors:** Thomas E. Wales, Aleksandra Pajak, Alžběta Roeselová, Santosh Shivakumaraswamy, Steven Howell, F. Ulrich Hartl, John R. Engen, David Balchin

## Abstract

The cellular environment is critical for efficient protein maturation, but how proteins fold during biogenesis remains poorly understood. We used hydrogen/deuterium exchange (HDX) mass spectrometry (MS) to define, at peptide resolution, the cotranslational chaperone-assisted folding pathway of *Escherichia coli* dihydrofolate reductase. On the ribosome, the nascent polypeptide folds via structured intermediates not populated during refolding from denaturant. Association with the ribosome allows these intermediates to form, as otherwise destabilizing C-terminal sequences remain confined in the ribosome exit tunnel. We find that partially-folded nascent chains recruit the chaperone Trigger factor, which uses a large composite hydrophobic/hydrophilic interface to engage folding intermediates without disrupting their structure. In addition, we comprehensively mapped dynamic interactions between the nascent chain and ribosomal proteins, tracing the path of the emerging polypeptide during synthesis. Our work provides a high-resolution description of *de novo* protein folding dynamics, thereby revealing new mechanisms by which cellular factors shape the conformational search for the native state.

## Introduction

The physical principles that underlie the refolding of small proteins in isolation are increasingly well understood (*1*). *De novo* protein folding *in vivo* differs, however, in that it begins in the context of translation. Unlike folding from denaturant, folding on the ribosome is coupled to vectorial synthesis (proceeding from the N to C terminus), such that not all elements of the nascent protein chain (NC) are simultaneously available for folding. As a result, structural motifs or domains may form sequentially during translation (*2–4*), a mechanism that can mitigate interdomain misfolding (*5–7*). The ribosome also places physical constraints on folding. The exit tunnel, which is narrow and negatively charged, limits the conformational space accessible to the nascent polypeptide (*8–10*), and proximity to the ribosome surface can destabilize complete domains (*11–15*). How these central features of cotranslational folding influence the resulting maturation pathway of the nascent polypeptide is incompletely understood.

The ribosome does not act alone in protein biogenesis, but associates with additional factors that regulate folding. Most prominent among these in bacteria is the highly abundant ribosome-bound chaperone Trigger factor (TF), which engages the majority of nascent *E. coli* proteins (*16*). Although not essential under normal growth conditions, TF deletion is lethal in the absence of buffering by the Hsp70 chaperone DnaK (*17*). As the paradigmatic cotranslational chaperone, various sometimes paradoxical functions have been ascribed to TF. These include inhibiting aggregation (*18*), promoting folding and assembly (*19, 20*), delaying cotranslational assembly (*21*), destabilizing folded domains (*22*), and favoring posttranslational folding (*23, 24*). How exactly chaperones modify *de novo* folding on the ribosome is unclear.

The inherent structural heterogeneity of the nascent protein, especially in the context of the size and complexity of the ribosome (with associated chaperones), poses a substantial technical challenge to probing local NC conformation. As a result, folding pathways on the ribosome are ill-defined, and whether translation fundamentally modifies the mechanism of protein maturation remains controversial (*25–27*). Here, we use a combination of proteomics, biophysical measurements and hydrogen/deuterium exchange (HDX) mass spectrometry (MS) to study cotranslational protein folding at the peptide level. Our approach resolves subtle local conformational differences between ribosome-bound folding intermediates, while simultaneously reporting on the dynamic behavior and interactions of ribosomal proteins and bound chaperones.

Using dihydrofolate reductase (DHFR) as a model, we show that the ribosome grants the NC access to an efficient folding route that is inaccessible during refolding from denaturant. Although the vectorial character of protein synthesis prevents simultaneous folding of all elements of the DHFR β-sheet, a central subdomain in DHFR behaves as a novel independent folding unit during translation. This is enabled by association with the ribosome, and occurs while the NC is bound by the chaperone TF. As a result, the nascent polypeptide is poised to complete folding immediately upon emergence of the C-terminus from the exit tunnel. Together, our data reveal how the ribosome and TF collaborate to define the structural progression of a nascent protein.

### Preparation of cotranslational folding intermediates for HDX MS analysis

As a model for studying protein biogenesis we used *E. coli* DHFR, an essential oxidoreductase enzyme with close homologues in all domains of life (*28*). DHFR is a single-domain monomeric protein of 159 aa, with a central eight-stranded β-sheet and four flanking α-helices (Figure 1A,C). The β-sheet is discontinuous in sequence: strand β1 inserts between β5 and β6, and β5 inserts between β1 and β2. DHFR refolding from denaturant *in vitro* has previously been characterized in detail, providing a point of comparison for folding on the ribosome (*29*). *In vitro*, global collapse occurs on the microsecond timescale, and the order of strands in the β-sheet motif is established within 6 ms. Final folding to the native state occurs in seconds via up to four parallel pathways. Importantly, folding of DHFR is substantially faster than synthesis on the ribosome, which would require ~8-16 s (~10-20 codons translated per second (*30*)). Stalling the ribosome is therefore not expected to distort nascent chain (NC) conformational sampling that occurs during normal translation.

**Figure 1.**
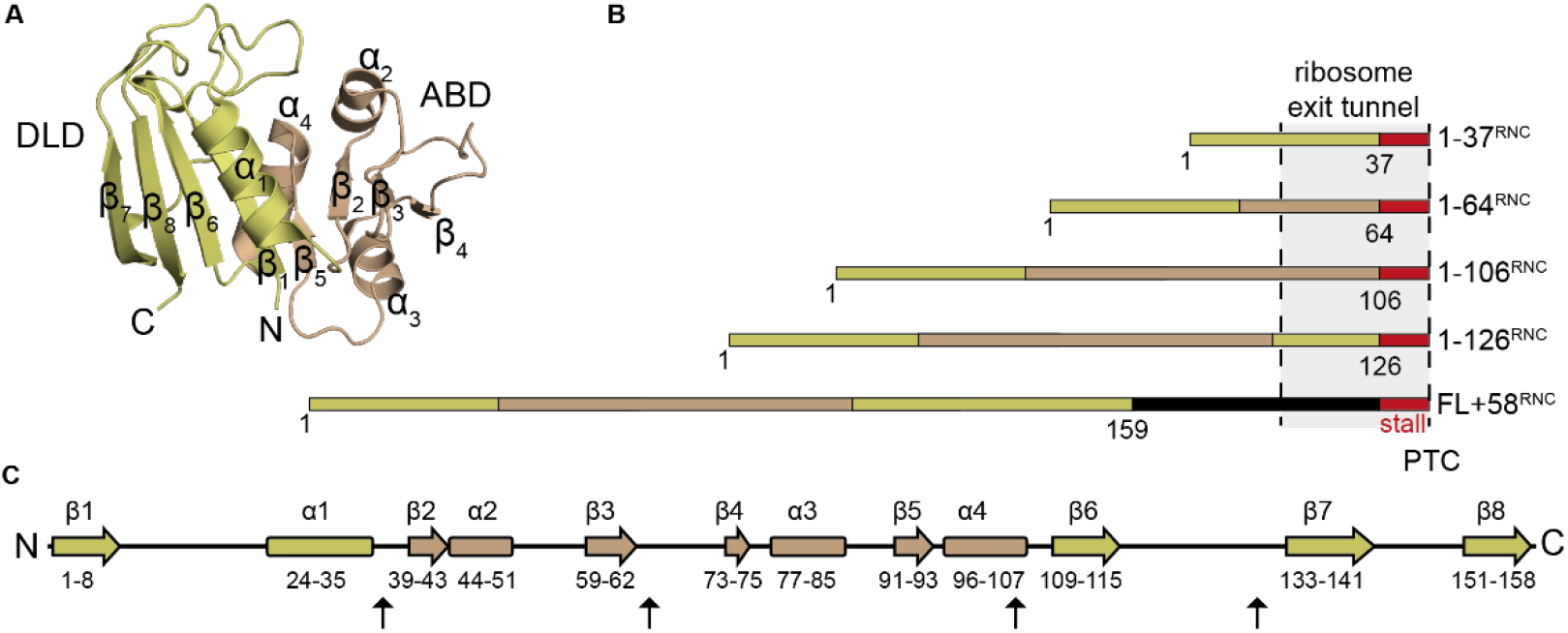
Design of DHFR ribosome:nascent chain complexes (RNCs). (**A**) Structure of *E. coli* DHFR (PDB: 5CCC). (**B**) Schematic illustrations of stall-inducing constructs. The stall site is indicated in red, and subdomains are colored gold (discontinuous loop subdomain, DLD) or bronze (adenosine binding subdomain, ABD), with the artificial linker in FL+58^RNC^ in black. The region of the nascent chain expected to span the exit tunnel is indicated. PTC, peptidyl transfer center. (**C**) Secondary structure elements in DHFR, colored according to subdomains as in (B). Sites where the stall sequence was inserted are marked with arrows.

To sample DHFR cotranslational folding intermediates along the pathway of vectorial synthesis, we prepared stalled ribosome:nascent chain complexes (RNCs) representing snapshots of folding *in vivo* (Figure 1B) (*31, 32*). The NC sequence was truncated at the boundaries between discrete structural motifs, in addition considering the annotation of DHFR “subdomains” (adenosine binding subdomain, ABD (residues 38-106); discontinuous loop subdomain, DLD (residues 1-37 and 107-159)). Translation was stalled at specific positions by encoding, C-terminal to each DHFR fragment, an 8 aa ribosome stall-inducing sequence derived from *M. succiniciproducens* SecM that is resistant to folding-induced release (*33*). As a control, we also prepared a construct consisting of the complete DHFR sequence separated from the stall site by an unstructured C-terminal linker of 50 aa, such that full-length DHFR is a total of 58 residues from the peptidyl transfer center (denoted FL+58^RNC^). Prior force profile analyses suggested that DHFR stalled at this linker length can fold into a conformation capable of binding the inhibitor methotrexate (*34*). Note that ~30 residues in an extended conformation are required to span the exit tunnel (*35*). The stall-inducing constructs were expressed in *E. coli* and RNCs were purified via an N-terminal affinity tag on the NC that was later cleaved (Figure S1A,B). The resulting RNCs were homogeneous, stable during prolonged incubation and over multiple freeze-thaw cycles, and insensitive to puromycin, indicating that the ribosomes were accurately stalled at the SecM sequence (Figure S1C-F). Quantitative label-free MS analysis confirmed the presence of the NC and all ribosomal proteins at close to the expected stoichiometries, and the absence of significant contamination (Figure S2A,B). The exception was the chaperone TF, which copurified predominantly with RNCs containing the first 106 or 126 residues of DHFR (1-106^RNC^ and 1-126^RNC^, Figure S1C and S2C,D).

With purified RNCs in hand, we could proceed with investigating the conformation of the NC, ribosome, and any associated proteins (e.g. TF). Purified RNCs were exposed to deuterated buffer for 10 s, 100 s or 1000 s and the entire system digested into peptides. The peptide mixture was separated by liquid chromatography and ion mobility, and the relative deuterium uptake of peptides from all proteins in the system was measured by mass spectrometry (Figure 2A and Data S1). The degree of deuterium incorporation reports on protein conformation, as backbone hydrogens are protected from exchange when involved in stable secondary structure, buried in the core of a folded protein or at a protein-protein interface (*36*). Comparative HDX measurements are therefore a sensitive probe of the local environment of amide hydrogens, and as such can readily distinguish folded versus unfolded and native versus non-native states, at the peptide level (*37–40*).

**Figure 2.**
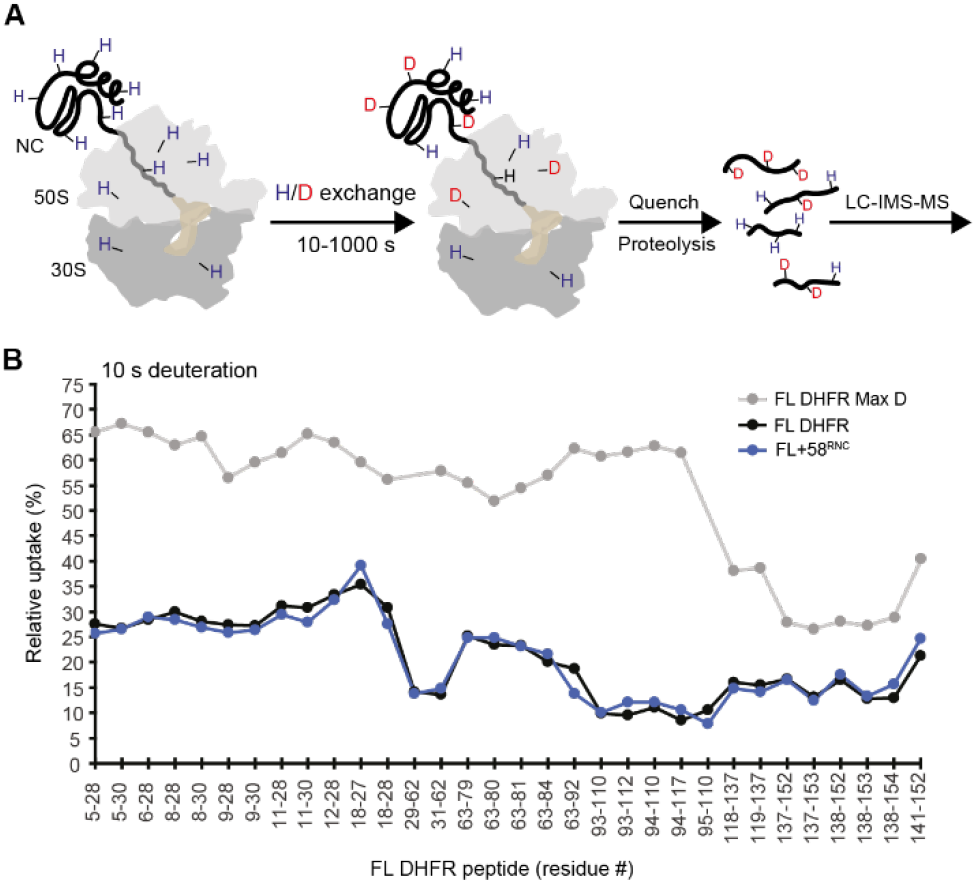
Hydrogen/deuterium exchange (HDX) mass spectrometry (MS) of RNCs. (**A**) Schematic of HDX MS experiment. Purified RNCs are deuterated for 10 s, 100 s or 1000 s, before the reaction is quenched and the entire system digested into peptides. Peptides are resolved in solution (LC, liquid chromatography) and in the gas phase (IMS, ion-mobility spectrometry), followed by analysis of deuterium incorporation by MS. (**B**) Relative deuterium uptake of DHFR peptides after 10 s exposure to deuterium, as a percentage of the maximum possible exchange, for isolated DHFR (FL DHFR) and FL+58^RNC^. A maximally deuterated control sample (FL DHFR Max D) is shown as a reference. Values are the average of 2-4 replicates. See also Figure S3 and Data S1.

To validate our methodology, we first analyzed full-length DHFR on the ribosome (FL+58^RNC^), where DHFR was expected to be completely emerged from the ribosome and folded. Thirty-four peptides from the NC covering 94% of the sequence could be followed, despite the analytical complexity of the system (Figure S3A). For comparison we analyzed isolated DHFR (here called FL DHFR), purified from *E. coli* as a natively folded, unliganded and untagged protein. Deuterium uptake for DHFR peptides was almost identical between FL+58^RNC^ and FL DHFR (Figure 2B, S3B and Data S1), indicating that DHFR, fully translated yet still coupled to the ribosome, is essentially natively folded. Two exceptions were subtle protection from exchange of peptide 63-92 in the NC, and deprotection of peptide 94-117, both apparent at longer deuterium exposure times. We hypothesize that these differences arise from weak interactions between the ribosome surface and folded DHFR, discussed in more detail below. Overall, these data demonstrate that our approach allows for analysis of HDX in an extremely complex mixture, and can accurately report on local NC conformation even in the background of the entire ribosomal protein complement.

### Cotranslational folding pathway of DHFR

To define the folding dynamics of DHFR on the ribosome we compared peptide-resolved HDX at different NC lengths (see Fig. 1B). FL DHFR and FL+58^RNC^ served as fully folded references, and deuterium uptake measured at 3 exchange times provided a fingerprint for the native conformation. We initially focused on the N-terminal region including strand β_1_ in the DLD (residues 5-30), which presented a set of overlapping peptides common to all RNCs. As a representative example, deuteration of peptide 9-28 as a function of NC length and deuterium exposure time is shown in Figure 3A. Note that the same behavior was observed in multiple peptides covering this region of DHFR (Figure 3B). At short chain lengths (1-37^RNC^), the N-terminal region showed intermediate levels of exchange, readily distinguishable from the same sequence in native DHFR which was much more protected at 10 s deuteration. The minor protection at short chain lengths may be due to interactions with the ribosome, or reflect transient non-native structure stabilized by the exit tunnel (*4*). Although substantially deprotected relative to native DHFR, the N-terminus was not completely deuterated in the RNCs, as judged by comparison to deuteration of a synthetic peptide comprising residues 1-37 (1-37^peptide^), which was maximally exchanged at the earliest labeling time. Elongation of the NC to extend the N-terminus fully beyond the exit tunnel (1-64^RNC^) resulted in further deuteration. The N-terminal region remained highly deuterated with little variation in uptake, even as the NC extended to 106 and 126 residues during synthesis of the ABD. Folding of the DLD is therefore delayed until the complete subdomain is available and the C-terminus is released from the ribosome, or artificially extended beyond the tunnel via a linker as in FL+58^RNC^. Notably, this folding pathway is different to the *in vitro* pathway of refolding of DHFR from denaturant, where the central 8-stranded β-sheet that spans the two subdomains is established early in the folding pathway (*41*).

**Figure 3.**
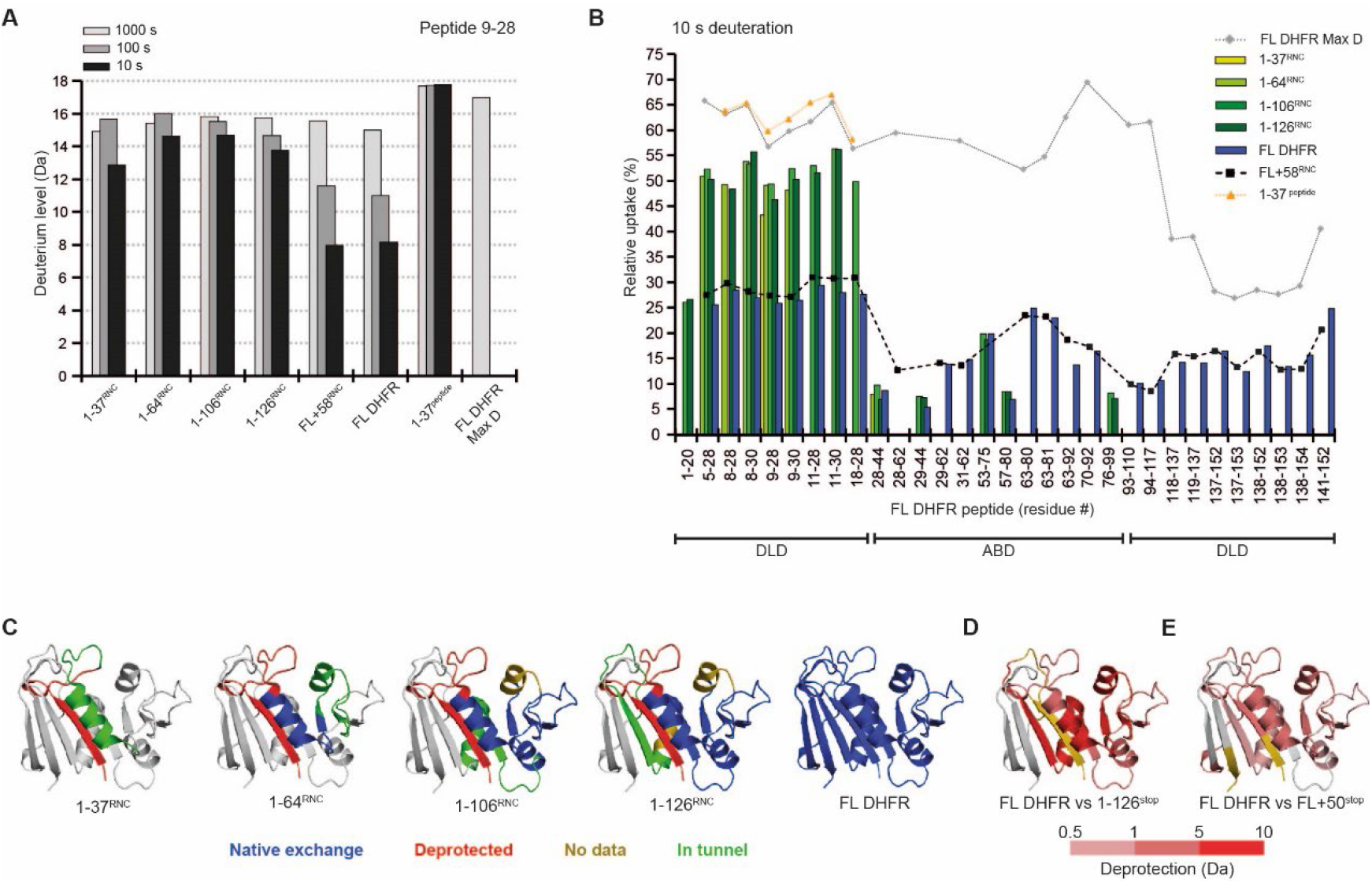
Conformational dynamics of nascent DHFR on the ribosome. (**A**) Deuterium incorporation into peptide 9-28 of DHFR as a function of deuterium exposure time and NC length. (**B**) Relative deuterium uptake of DHFR peptides after 10 s exposure to deuterium. Peptides belonging to the DLD or ABD subdomains are indicated. Values are the average of 2-4 replicates. (**C**) Peptide-resolved folding of nascent DHFR, illustrated on the structure of native DHFR (PDB: 5CCC). Peptides with the same deuterium uptake (within 0.5 Da) as the native controls (FL DHFR or FL+58^RNC^) are colored blue, while peptides that are deprotected relative to the controls are colored red. Regions for which we did not measure HDX are colored gold. The C-terminal 22 aa in each construct, expected to be occluded in the exit tunnel at that chain length, is colored green. The part of DHFR that is not yet synthesized at each chain length is colored white. (**D**) Difference in deuterium uptake between FL DHFR and 1-126^stop^ after 10 s deuteration. Darker red indicates more deuterium in 1-126^stop^ relative to FL DHFR (increasing deprotection). Residues 127-159, not present in 1-126^stop^, are colored white. (**E**) Difference in deuterium uptake between FL DHFR and FL+50^stop^ after 100 s deuteration. Darker red indicates increasing deprotection of FL+50^stop^ relative to FL DHFR. Peptides that do not differ in exchange are colored white. See also Figure S4 and Data S1.

Extending this analysis to additional peptides across the RNCs allowed us to reconstruct the complete cotranslational folding pathway of DHFR. Peptide-resolved HDX for each different length RNC is shown in Figure 3B, S4A,B and Data S1, and folding information is mapped onto the structure of native DHFR in Figure 3C. Although not all peptides were uniformly detected across stalled RNCs of different length, clear patterns of HDX protection corresponding to NC folding were apparent. As described above, the N-terminal strand β1 is initially unfolded (1-37^RNC^) when exposed outside the exit tunnel. In 1-64^RNC^, residues 25-36 emerge from the tunnel and fold into a native helix α1 belonging to the DLD, while β1 remains unstructured. Synthesis of the remainder of the ABD (1-106^RNC^) allows β2-4 to exit the ribosome and acquire native-like protection from HDX. This protection was not due to association with TF, as discussed in subsequent sections. Folding of the ABD completes when 126 residues of DHFR have been synthesized (1-126^RNC^), allowing β5 and α4 to coalesce with the remainder of the ABD outside the exit tunnel. At this point β6 is still occluded in the tunnel, precluding folding of the adjacent β1 in the DLD. Synthesis and release of the C-terminal strand β8 from the ribosome triggers final folding of the DLD including β1 (FL DHFR and FL+58^RNC^).

Our detailed analysis of cotranslational folding revealed a strong propensity of nascent DHFR to adopt partially structured states on the ribosome. We therefore considered whether structure prediction methods could discern patterns of folding in nascent polypeptides containing incomplete sequence information. We used the Colabfold implementation of Alphafold2 (*42, 43*) to predict the structures of truncated DHFR chains corresponding to intermediates in vectorial synthesis (Figure S5). The predicted structures for the DHFR fragments corresponding to 1-64^RNC^ and 1-106^RNC^ are in qualitative agreement with our experimental results (Figure S5). Specifically, the models recapitulate our observation that the ABD can begin to fold while the N-terminus remains unstructured (Figure 3B,C), offering independent support for our conclusion that the ABD can constitute an independent folding unit. However, there are clear deviations between our measurements and the predictions from Alphafold2 at short and long chain lengths. In 1-37^RNC^, the nascent polypeptide is incorrectly modelled as a stable helix, while in 1-126^RNC^ the N-terminal strand β1 is interpreted to be folded, inconsistent with our data showing that this region only folds when the DLD is completely synthesized. These results emphasize that folding intermediates, even when maintained at equilibrium as in our stalled RNCs, are not well modelled by current native-state-centric approaches.

### Association with the ribosome stabilizes a cotranslational folding intermediate

We noticed that unlike the DLD, peptides corresponding to the β-sheet of the ABD became progressively protected from HDX during translation, requiring neither the sequence context of full-length DHFR nor even the complete subdomain (Figure 3C). In contrast, previous work showed that fragments of DHFR produced by chemical cleavage are disordered in isolation (*44*). To test whether the ribosome modulates the conformation of the nascent ABD, we attempted to express isolated fragments corresponding to the ABD (1-64, 1-106 and 1-126) in *E. coli.* Only the longest fragment consisting of residues 1-126 (herein 1-126^stop^) was soluble, and this was contingent on maintaining it as a fusion protein, with monomeric ultrastable GFP (muGFP (*45*)) at the N-terminus. We analyzed 1-126^stop^ using HDX MS, which revealed substantial deprotection relative to native DHFR (Figure 3D and Data S1). High levels of exchange were observed for the N-terminal region, as expected in the absence of the C-terminal strands β7 and β8 that complete the DLD. The β-strands (β2-4) and peripheral helices comprising the core of the ABD were also strongly deprotected relative to full-length DHFR. The ABD therefore fails to fold into a stable structure in isolation, although native-like folding of this subdomain is supported on the ribosome in the context of an RNC.

Why can incomplete chains of DHFR fold on the ribosome but not in free solution? We noticed that residues 93-118 near the C-terminus of 1-126^stop^ were maximally deuterated even at short deuteration times, indicative of structural disorder (Figure 3D and Data S1). In the corresponding RNC, however, this sequence is sterically confined in the ribosome exit tunnel (Figure 1B and 3C). We therefore considered whether, in the absence of the ribosome, a region of C-terminal disorder may destabilize folded DHFR. Indeed, previous work has shown that unstructured termini can generate an entropic force that modulates protein conformation (*46*). To explore this idea, we used FL+58^RNC^ as a model. FL+58^RNC^ consists of full-length DHFR tethered to the PTC via a flexible 50 aa linker that is largely occluded in the exit tunnel (Figure 1B). Our HDX analysis showed that DHFR is natively folded in this context and almost indistinguishable from the isolated domain (Figure 2B and S3B,C). We hypothesized that releasing the protein from the ribosome would remove the protective effect of the exit tunnel and allow an unstructured terminus to disrupt the conformation of DHFR. To test this, we replaced the stalling sequence with a stop codon, resulting in FL+50^stop^ which was expressed in *E. coli* and purified as a soluble protein. HDX MS analysis of FL+50^stop^ confirmed that the linker was highly dynamic and easily deuterated, as expected (Figure S4C-E and Data S1). Furthermore, we observed extensive deprotection relative to native FL DHFR, consistent with increased structural dynamics (Figure 3E and S4C-E). Deprotection was strongest in peptides covering β1-3, α1 and α2, indicating that neither the DLD nor ABD is completely folded in FL+50^stop^. Combined, our observations suggest a model whereby domain instability induced by unstructured C-termini is rescued by confinement in the ribosome exit tunnel, potentially facilitating the folding of structured intermediates during translation.

### DHFR can fold to a hyperactive state close to the ribosome and unimpaired by TF

Close proximity to the ribosome surface was previously shown to destabilize some nascent proteins (*11–13, 15*). However, our sensitive peptide-resolved HDX analysis of FL+58^RNC^ did not reveal substantial differences in conformation compared to isolated DHFR (Figure 2B). As a more stringent test, we prepared an RNC with a 38-residue linker (FL+38^RNC^), designed to bring DHFR much closer to the ribosome surface. HDX MS showed that the folded core of DHFR in FL+38^RNC^ was similar in protection to isolated DHFR (Figure 4A, S6A and Data S1). However, observe clear differences in peripheral loops and part of the DLD. Peptides covering residues 5-28 and 63-92 were protected from HDX in FL+38^RNC^, while peptide 94-117 was deprotected. We also noted that a minor fraction (<5%) of the FL+38^RNC^ population copurified with TF (Figure S2C). As an orthogonal test of folding, we measured the DHFR enzyme activity of the purified RNCs (Figure 4B). Despite being coupled to ribosomes, both FL+58^RNC^ and FL+38^RNC^ were enzymatically active, consistent with our HDX results showing that the NC is essentially natively folded. Close proximity to the vestibule of the ribosome exit tunnel does not, therefore, substantially disrupt the conformation of folded DHFR.

**Figure 4.**
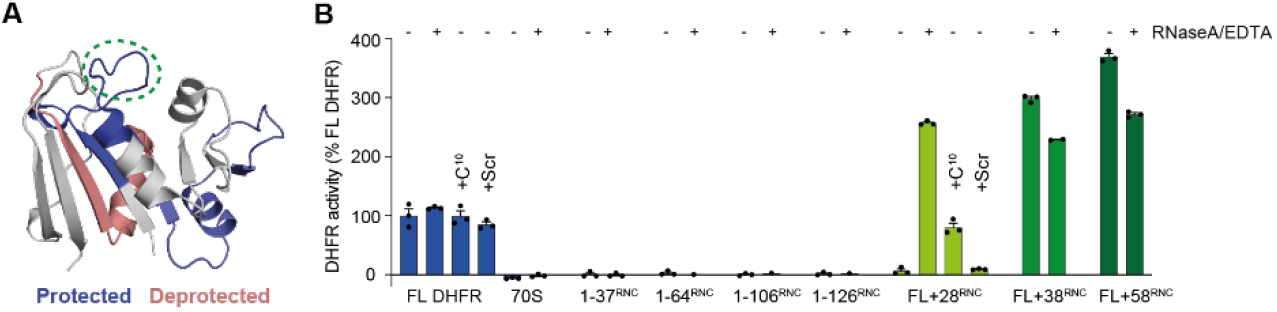
Ribosome contacts modulate the structure of nascent DHFR without inducing unfolding. (**A**) Relative deuterium uptake of FL+38^RNC^ compared to FL DHFR. DHFR peptides that are protected in the RNC relative to FL DHFR, at any deuteration time point, are colored blue, deprotected peptides are colored red. The Met20 loop is circled. See also Figure S6A and Data S1. (**B**) Oxidoreductase activity of 25 nM FL DHFR, empty 70 S ribosomes without a NC, or RNCs. Where indicated, reactions are supplemented with 50 μg/mL RNaseA and 50 mM EDTA, a peptide corresponding to the C-terminus of DHFR (C^10^, SYCFEILERR), or a scrambled-sequence control peptide (Scr, RFIERCELYS).

Inspection of the enzyme activity data revealed, remarkably, that FL+38^RNC^ and FL+58^RNC^ were at least 3-4 fold more active than isolated DHFR, with increased *V*_max_ and *K*_M_^DHF^ (Figure 4B and Figure S6C). The extent of activity stimulation may be underestimated, because RNC concentration was calculated by assuming 100% occupancy of NCs on ribosomes (See Methods). This was *bona fide* DHFR activity, as it was fully inhibited by methotrexate, and neither empty ribosomes nor the intermediate length RNCs showed detectable activity (Figure 4B and S6D). We observed the same effect for an RNC with a different linker sequence, indicating that hyperactivity is not an artefact of linker chemistry (Figure S6B). Notably, the region near the N-terminus of DHFR that is protected in FL+38^RNC^ includes the “Met20 loop” at the folate binding site of DHFR (Figure 4A), the dynamics of which are known to strongly influence catalysis (*28*). These data imply that physical interactions with the ribosome alter the active site of DHFR without inducing global unfolding.

To test the threshold linker length for folding of DHFR on the ribosome, we prepared an RNC with the spacer between the PTC and C-terminus shortened to 28 aa (FL+28^RNC^). This RNC was devoid of oxidoreductase activity (Figure 4B) and copurified with TF (Figure S2C). TF was not responsible for the lack of activity, because FL+28^RNC^ purified from a TF-free background was also enzymatically inactive (Figure S6B). We hypothesized that a 28 aa linker is insufficient to expose the C-terminus of DHFR outside the exit tunnel, precluding folding of the DLD. Indeed, DHFR activity could be restored by releasing the NC from FL+28^RNC^ with EDTA/RNase, which disrupts ribosomes and cleaves the peptidyl tRNA (Figure 4B). This observation is consistent with our interpretation, based on HDX MS, that DHFR folding completes post-translationally when the C-terminus emerges from the tunnel (Figure 3A-C). To test whether DHFR in FL+28^RNC^ is folding-competent prior to release from the ribosome, we added a peptide comprising the C-terminal 10 aa of DHFR (peptide C^10^) in *trans.* Peptide C^10^ conferred enzyme activity to FL+28^RNC^ in a concentration-dependent manner, whereas a scrambled-sequence control had no effect (Figure 4B and S6B). None of the peptides influenced the activity of isolated full-length DHFR. TF was not required for reactivation, since TF-free FL+28^RNC^ was similarly reactivated (Figure S6B). Furthermore, pelleting experiments showed that addition of peptide C^10^ did not induce release of the NC from ribosomes, nor did it displace TF (Figure S6E). DHFR is therefore poised to complete folding upon emergence of the C-terminus from the ribosome exit tunnel, and neither close proximity to the ribosome surface nor association with TF prevent acquisition of the native state.

### Interaction of Trigger factor with DHFR on the ribosome

We next sought to determine the role of Trigger factor in cotranslational folding of DHFR. TF is a 48 kDa ribosome-associated chaperone with a three-domain architecture (Figure 5A). The ribosome binding domain (RBD) contains a conserved ribosome-interaction motif (*47*), and the substrate binding domain (SBD) is required for chaperone function off the ribosome (*48*). Although the peptidyl-prolyl isomerase domain (PPD) catalyzes *cis*/*trans* proline isomerization in isolated proteins *in vitro*, its role in *de novo* folding on the ribosome is enigmatic (*49*). Due to the highly dynamic nature of the chaperone:RNC complex, details of the interaction between TF and NCs have eluded structural characterization (*50, 51*), although site-specific photocrosslinking showed that NCs contact all three domains of TF (*51, 52*).

**Figure 5.**
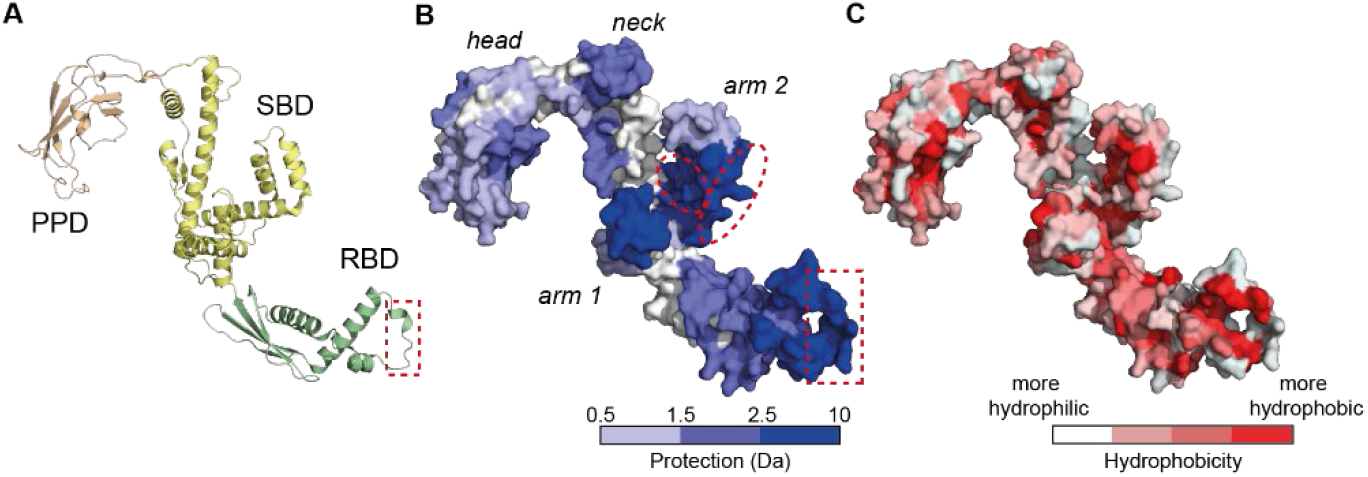
NC interactions with Trigger factor (TF). (**A**) Structure of *E. coli* TF (PDB: 1W26). RBD, ribosome binding domain; SBD, substrate binding domain; PPD, peptidyl-prolyl isomerase domain. The ribosome binding motif in the RBD is boxed. (**B**) Difference in deuterium uptake between isolated TF and TF bound to DHFR RNCs after 10 s deuteration. Darker blue indicates less deuteration of RNC-bound TF relative to isolated TF (increasing protection). The sites on arm 2 that preferentially engages 1-126^RNC^ are marked by ellipses. See also Data S1. (**C**) TF surface colored according to hydrophobicity (*83*).

TF copurified predominantly with 1-106^RNC^ and 1-126^RNC^, coinciding with synthesis of the complete ABD (Figure S1C and S2C). To characterize the interaction between TF and nascent DHFR, we analyzed deuterium uptake for endogenous TF in the RNC samples and compared these data to HDX MS measurements of isolated TF. In total we followed 181 peptides for TF in each condition, with 99.5% sequence coverage (Figure S7A and Data S1). A potential confounding factor for this analysis is that isolated TF is weakly dimeric (dimerization Kd ~1 μM), but binds ribosomes as a monomer via interfaces that are partially occluded in the dimer (*52–54*). To account for this dimerization effect, we used both wild-type TF and a constitutively monomeric variant (*55*) as reference controls for isolated TF. HDX MS analyses confirmed that the monomeric variant was deprotected, relative to isolated wild-type TF, at sites that are normally buried by the dimer interface (Figure S7B). We found that the same regions of TF were protected when RNC-associated TF was compared to either isolated monomeric TF or wild-type TF (Data S1). The amount of protection was greater when monomeric TF was the reference, consistent with the reported competition between TF dimerization and ribosome binding (*53*).

Compared to isolated TF, TF that was bound to 1-106^RNC^ and 1-126^RNC^ was very strongly protected from HDX in the ribosome-interacting motif of the RBD, as expected (*47*) (Figure 5B). We also observed protection of additional regions in all three domains of TF, which we attribute to interaction with nascent DHFR (Figure 5B). Strong protection was observed in the two “arms” of the SBD, with intermediate protection in the RBD, “neck” region and catalytic site in the PPD. Analysis of the hydrophobicity of the binding interface revealed a mix of hydrophobic and hydrophilic surfaces (Figure 5C). The interaction sites in the RBD are predominantly hydrophobic, whereas hydrophilic surfaces in the PPD are preferred. In the SBD, the interface includes the hydrophobic pocket in the crook of arm 2, as well as hydrophilic surfaces in arm 1 and the neck. The hydrophilic part of the neck situated between the arms, previously implicated in substrate engagement off the ribosome (*56*), was not protected from exchange. The protected sites were the same in TF bound to 1-106^RNC^ and 1-126^RNC^, indicating a common interaction surface. We did, however, detect increased protection of hydrophilic regions in arm 2 (residues 351-374 and 388-409) when TF was bound to the longer NC (Figure 5B, red ellipses, and Data S1).

### Nascent DHFR can fold while associated with Trigger factor

To determine how TF influences the folding of DHFR on the ribosome, we purified 1-106^RNC^ and 1-126^RNC^ from *E. coli* lacking TF (*17*), resulting in 1-106^RNCΔTF^ and 1-126^RNCΔTF^. MS analysis confirmed that the absence of TF did not result in other chaperones (e.g. DnaK or DnaJ) copurifing with the RNCs (Figure S2C). Furthermore, pelleting assays showed that 1-126^RNCΔTF^ was still competent to bind purified TF *in vitro*, and binding was sensitive to mutation of the ribosome-interacting motif in TF (Figure S8A).

To probe the conformation of the nascent chain without TF, we analyzed the HDX behavior of nascent DHFR in 1-106^RNCΔTF^ and 1-126^RNCΔTF^ (Figure S7C and Data S1). A set of peptides reported on the N-terminus and β-strands in the ABD, enabling us to compare WT and ΔTF RNCs. Although some marginal differences in deuterium uptake could be detected in the absence of TF, these were not consistently observed across overlapping peptides. Cotranslational folding of the ABD therefore occurs irrespective of the presence of TF, and TF binding does not explain our observation that the N-terminus remains unfolded until release of the C-terminus from the ribosome (Figure 3C).

Considering that DHFR folding can occur in the presence of TF, what features of the NC are recognized by the chaperone? Our data show that TF binding coincides with conformations of nascent DHFR characterized by a folded ABD and unstructured DLD (Figure 3C), and that binding occurs via a mixed hydrophobic/hydrophilic interface (Figure 5B). To probe the contribution of electrostatic versus hydrophobic interactions to binding, we tested the salt-sensitivity of the TF:RNC interaction. Quantitative proteomic analysis revealed a preference of TF for 1-106^RNC^ over 1-126^RNC^ when the complexes were purified under high-salt conditions (1 M KOAc), with ~2-fold higher occupancy of the chaperone on the shorter NC (Figure S2C). In contrast, RNCs purified under low-salt conditions (100 mM KOAc) bound similar amounts of TF, such that TF was stoichiometric to ribosomes in both cases (Figure S8B). TF binding to 1-126^RNC^ is therefore partially stabilized by electrostatic interactions, unlike binding to 1-106^RNC^ which is predominantly hydrophobic and therefore stabilized by high salt. This observation is consistent with folding-induced burial of the ABD hydrophobic core in 1-126^RNC^ (Figure 3C), as well as the preference of this NC for binding hydrophilic surfaces on TF (Figure 5).

To directly test the contribution of NC folding to TF binding, we simultaneously introduced three mutations (V40A, V75H, I91L) into 1-126^RNC^, designed to destabilize the native ABD (*57*) (Figure S8C). We found that the mutated RNC bound more TF than WT 1-126^RNC^ when purified under high-salt conditions (Figure S8D). This difference was less pronounced, however, when binding was reconstituted *in vitro* under low-salt (Figure S8E). Binding of TF to the destabilized RNC is therefore less dependent on electrostatic interactions, suggesting that TF engages poorly folded intermediates via hydrophobic surfaces. Taken together, these observations indicate that TF uses a composite hydrophobic/hydrophilic interface to accommodate both folded and unfolded NCs, and provide indirect evidence supporting our conclusion that the ABD is natively folded in WT 1-126^RNC^. Our low-salt conditions are in a similar range of ionic strength to the *E. coli* cytosol (~100-200 mM (*58*)). *In vivo*, TF would therefore be expected to bind equally well to NCs exposing different amounts of hydrophobic surface, exploiting different types of interaction in each case.

### Nascent chain length-dependent interactions with ribosomal proteins

We next considered how nascent DHFR might influence ribosomal proteins and their conformation, with the goal of identifying possible NC:ribosome interaction sites. To this end, we compared the HDX of ribosomal proteins in the RNCs to the same proteins in empty 70S ribosomes. We focused our analysis on a set of five ribosomal proteins (L4, L22, L23, L24 and L29) that are near/in the exit tunnel and therefore likely to contact the emerging NC (Figure 6A). Sequence coverage of these ribosomal proteins was close to 100% (Figure S9A). We identified several sites of HDX protection in ribosomal proteins when NC was present, often dependent on NC length (Figure 6B-G and Data S1).

**Figure 6.**
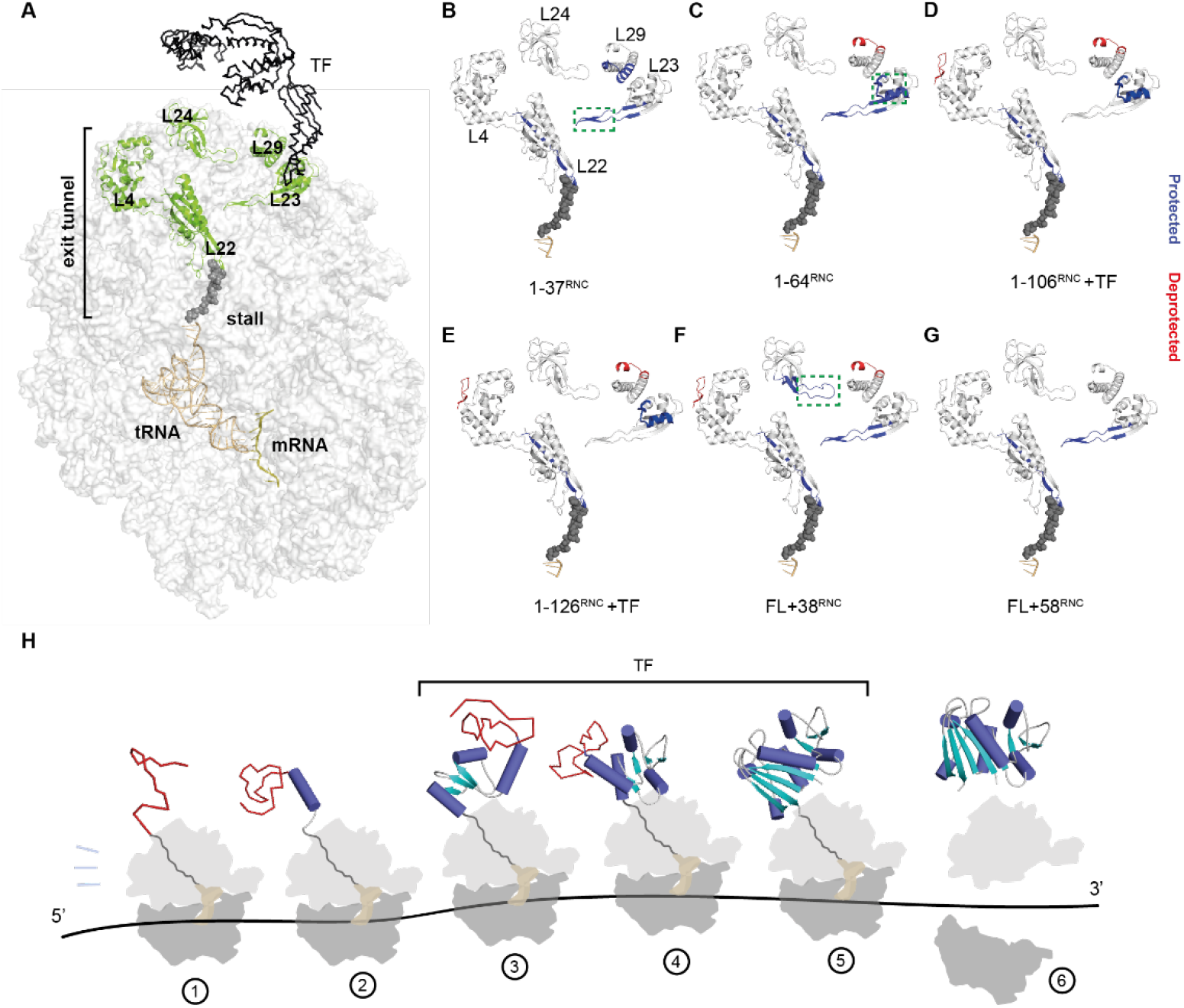
Effect of nascent DHFR on ribosomal proteins. (**A**) Side view of the 70S ribosome, highlighting ribosomal proteins that line the exit tunnel and vestibule (PDB: 3JBU). TF is placed onto the structure based on (*51*). The structure contains the 17 aa wild-type SecM stall-inducing sequence, although to match the length of the sequence used in this study only 8 aa are shown. (**B-G**) HDX analysis of ribosomal proteins in RNCs. Peptides that are protected from HDX in each RNC relative to empty ribosomes, at any deuteration time point, are colored blue. Deprotected peptides are colored red. The tunnel-facing loop of L23 is boxed in **B**, the TF docking site is boxed in **C**, and the exposed loop of L24 is boxed in **D**. See also Data S1. (**H**) Schematic biogenesis pathway of DHFR, based on HDX MS analysis of stalled RNCs. Non-native elements are colored red. The N-terminus of DHFR emerges from the ribosome in an unstructured state and interacts weakly with L29 at the exit port (1). Synthesis of an additional 27 residues results in folding of helix α1 and exposes a basic patch that contacts the surface-exposed loop of L23 (2). The remainder of the adenosine binding subdomain folds cotranslationally and while engaged by TF docked at L23 (3,4). The nascent subdomain is protected from denaturation by occlusion of the succeeding sequence in the exit tunnel. Folding of the DLD, including the N-terminal strand, occurs when the C-terminus is provided as a peptide in *trans* (5) or when the complete domain is released from the exit tunnel upon termination of synthesis (6).

In our RNCs, the lower part of the tunnel is occupied by the stall-inducing sequence. Consequently, the tunnel-exposed loop of L22, which together with L4 forms the constriction site close to the PTC, was protected from exchange at all NC lengths. The corresponding loop in L4 was not protected, potentially because the stall sequence interacts less stably with this site. Consistent with this interpretation, structural data show that wild-type SecM is positioned closer to L22 than L4 (*59*)(Figure 6A). Peptide 22-42 of L29, located on the ribosome surface adjacent to the exit port, was protected only in the shortest RNC (1-37^RNC^) (Figure 6B). Longer NCs may expose sequences that favor different interaction sites (discussed below) or preferentially interact with TF rather than L29.

Peptide 8-29 of L23, constituting the binding site for TF on the ribosome surface (*47*), was strongly protected in 1-106^RNC^ and 1-126^RNC^ (Figure 6D,E), as expected since these RNCs recruit TF (Figure S2C). However, we also detected protection of the TF docking site in 1-64^RNC^ which does not engage TF (Figure 6C). This protection is presumably due to interaction with the NC, and is consistent with the sequence characteristics of nascent DHFR. In 1-64^RNC^ (but not 1-37^RNC^), the emerging NC is highly basic (pI ~9.9), perhaps facilitating electrostatic interactions with the acidic TF-docking site. Direct competition between the NC and the TF-binding site on the ribosome may constitute an additional mechanism by which TF recruitment to RNCs is regulated.

Peptide 61-84 of L23, forming a hairpin loop that protrudes into the exit tunnel ~50 Å from the PTC (Figure 6B, green box), was protected in all RNCs except 1-106^RNC^ and 1-126^RNC^. Interestingly, these are also the two RNCs that most strongly recruit TF (Figure S2C), suggesting the possibility of allosteric communication between the NC and TF via L23, as previously hypothesized (*60*). To probe this phenomenon further, we analyzed the HDX behavior of ribosomal proteins in 1-106^RNCΔTF^ and 1-126^RNCΔTF^ (Figure S9B,C and Data S1). Removing TF did not result in new protection of the L23 tunnel loop, indicating that TF does not reciprocally influence the conformation of L23. Rather, TF binding correlates with intrinsic features of the NC that determine tunnel interactions.

In FL+58^RNC^ none of the proteins surrounding the exit port were protected from HDX relative to empty ribosomes, consistent with the absence of stable interactions with the ribosome surface (Figure 6G). However, analysis of FL+38^RNC^ revealed protection of a loop in L24 (Figure 6F, green box), previously identified to interact with NCs (*13*). Interactions with L24 may underlie the subtle modulation of DHFR conformation that we observe for this RNC (Figure 4A).

In addition to protection from exchange relative to empty ribosomes, we noted several regions for which deprotection was induced by the NC (Figure 6B-F). Deprotection occurred in specific peripheral regions of L4 and L29, distinct from NC contact sites near the exit tunnel and port. Direct NC interactions with the ribosome could therefore allosterically influence the conformational dynamics of distant sites on the ribosome surface.

Together, our HDX MS analysis of ribosomal proteins suggests that nascent DHFR does not generically interact with proteins comprising the exit tunnel, but rather samples a biased route during synthesis that is potentially dictated by the chemical properties of the emerging sequence. Specific NC:ribosome interactions may further modulate the pathway of cotranslational folding (*10*).

## Discussion

We have presented a comprehensive analysis of the conformational dynamics of a ribosome:chaperone:nascent chain system trapped at various stages during protein synthesis. Unlike previous strategies for studying cotranslational folding (*61*), our HDX MS approach is label-free, yields local information (resolved to peptide level), and simultaneously reports on the structural dynamics of all proteins in the system. This has allowed us to directly compare the folding pathway of a model single-domain protein on and off the ribosome, as well as define the contribution of a ribosome-associated chaperone to folding.

We find that, in contrast to previous models of folding on the ribosome (*3, 13, 62*) cotranslational folding of DHFR is neither strictly sequential (N- to C-terminal) nor concerted (all-or-none). Rather, resolving *de novo* folding at peptide-level has revealed a combination of both mechanisms. Sequential folding of the middle subdomain poises DHFR for rapid completion of folding in a concerted post-translational step involving both the N- and C-termini (Figure 6H). This may represent a generic mechanism exploited by proteins like DHFR with discontinuous domain organizations, to minimize the delay between synthesis and acquisition of the native state.

Importantly, the efficient cotranslational folding pathway we observed for DHFR is not supported off the ribosome. During refolding *in* vitro, elements of the central β-sheet of DHFR fold simultaneously rather than sequentially, and intermediates with a folded ABD are not populated. Therefore, the ribosome chaperones nascent DHFR in at least two ways. First, by acting as a potent solubility tag (*63*), the ribosome allows the NC to access conformations that are too aggregation-prone to persist in isolation. Second, occlusion of the dynamic C-terminus in the exit tunnel prevents entropic destabilization of vulnerable folding intermediates. Disordered termini can alter the stability and dynamics of folded proteins via excluded volume effects (*46, 64*), a phenomenon that we reproduce for DHFR using an artificial unstructured extension. Here, we provide evidence that the ribosome buffers this effect during protein synthesis. In this regard, the physical dimensions of the exit tunnel, which are conserved in cytosolic ribosomes (*65*), may have facilitated the evolution of topologically complex folds. Recent work showed that the N-terminal domain of nascent elongation factor G is denatured by the succeeding unstructured domain (*7*). We speculate that entropic destabilization underlies the observed interference between domains, which is mitigated by the exit tunnel when the N-domain is close to the ribosome. Interestingly, destabilization of the upstream domain at longer linker lengths was prevented by TF (*7*). In this scenario TF may function to extend the exit tunnel, thereby protecting folded domains/intermediates from entropic destabilization by longer unstructured sequences. Other factors, such as interactions of the NC with the ribosome surface (Figure 6B-G), are also likely to play a role in modulating the stability of cotranslational folding intermediates (*10, 66*).

Although the chaperone function of TF off the ribosome is well characterized (*18, 19, 48, 56, 67*), its canonical role as a cotranslational chaperone is comparatively poorly understood. Indeed, little is known in general about how chaperone activity manifests at the ribosome (*68*). Here we address two key outstanding questions: 1) considering that TF binds the majority of nascent proteins, what properties of NCs are recognized? 2) How does TF engagement influence the conformation of the NC? We find that NCs with little compact structure fail to engage TF, as evidenced by the lack of binding to 1-37^RNC^ and 1-64^RNC^, as well as FL+58^RNC^ which presents an unstructured linker to the chaperone. This argues against the unstructured N-terminus being the driver of TF binding to intermediate length RNCs. Instead, our data indicate that TF prefers partially (1-106^RNC^, 1-126^RNC^) or fully-folded (complemented FL+28^RNC^) domains (Figure 6H). Native-like folding is not, however, required for TF binding, shown by the strong recruitment of TF to an RNC exposing a destabilized subdomain. We propose that this remarkable plasticity in binding is achieved via dynamic sampling of an unusually large chaperone:NC interface that includes both hydrophilic and hydrophobic surfaces (Figure 5), reconciling competing models for substrate recognition by TF (*18, 19*). In our model, persistent binding to RNCs would require simultaneous engagement of multiple sites across the TF surface, with ribosome binding by TF contributing to overall avidity. The chemically heterogeneous binding surface accommodates both partially and fully folded domains, allowing continuous engagement of NCs as they mature. Folding may therefore occur while the NC is associated with TF (*56*). The requirement for multiple distributed interaction sites would also explain why very short NCs are not stably engaged by TF (*16*), regardless of their folding status (Figure 6H). Importantly, the binding mode we describe allows TF to engage fragile folding intermediates without disrupting incipient structure. Indeed, we observe that cotranslational folding intermediates of DHFR are essentially identical regardless of the presence of TF. This mechanism may be particularly important for efficient folding of aggregation-prone NCs, which would be protected from aberrant interactions without reversing folding or preventing cotranslational assembly (*21*). TF is therefore mechanistically distinct from the Hsp70 chaperone system, which also engages nascent polypeptides (*17, 69*) but competes with client folding (*70*).

Previous work has shown that interactions with the ribosome surface destabilize the NC, thereby delaying folding relative to synthesis (*11, 12, 14, 15*). We find that this is not necessarily a general phenomenon. The conformational dynamics of the folded core of DHFR are unperturbed by proximity to the ribosome, and our peptide complementation experiments demonstrate that DHFR can fold to an enzymatically active state very close to the ribosome surface (Figure 6H). Surprisingly, rather than disrupting folding, subtle modulation of the DHFR active site at the ribosome stimulates the activity of the nascent enzyme. This observation raises the possibility that ribosome interactions may positively modulate the function of N-terminal domains in multidomain proteins, for example by promoting cofactor loading during synthesis.

Although we chose to analyze in detail only the NC and a small subset of ribosomal proteins that directly engage the NC, the high-quality HDX MS dataset we describe here is comprehensive. HDX MS data for all proteins in the system are included within our PRIDE archive database submission for future analysis. Our approach could shed new light on other aspects of translation regulation, especially dynamic processes that have been challenging to study using conventional structural biology methods. These include enzyme and chaperone binding to nascent chains, ribosome assembly, and ribosome dynamics during elongation.

## Acknowledgements

We thank John Christodoulou (University College London) for the kind gift of the *Δtig E. coli* strain; Svend Kjaer (Francis Crick institute) for GFP-DARPin affinity resin, 3C protease and TEV protease; Stephane Mouilleron (Francis Crick institute) for 3C protease and TEV protease, Keith Fadgen (Waters) for technical assistance with HDX MS, and the Peptide Chemistry Science Technology Platform at the Francis Crick institute for peptide synthesis.

## Authors contributions

Conceptualization: F.U.H., J.R.E., and D.B. Investigation: T.E.W., A.P., A.R., S.S., S.H., and D.B. Formal analysis, Data curation, and Visualization: T.E.W., A.P., S.H., J.R.E., and D.B. Writing – original draft: T.E.W., A.P., and D.B. Writing – review & editing: all authors.

## Data and materials availability

All mass spectrometry data have been deposited to the ProteomeXchange Consortium via the PRIDE partner repository with the dataset identifiers PXD036784 and PXD036945.

## Supplementary materials

Materials and Methods

Fig. S1 to S9

Table S1

Data S1

