## Supplemental information for "Resolving chaperone-assisted protein folding on the ribosome at the peptide level"

**This file includes:**

Materials and Methods

Figs. S1 to S9

Table S1

Caption for Data S1

**Other Supplementary Materials for this manuscript include the following:**

Data S1

### Materials and Methods

Expression and purification of ribosome-nascent chain complexes (RNCs)

Expression constructs were synthesized by Twist Biosciences (San Francisco, USA). All ORFs were cloned between a ribosome binding site (TTTGTTTAACTTTAAGAAGGAGA) and 6xHis-tag followed by a stop codon in pET21 plasmids with ampicillin resistance. The amino acid sequences encoded by each ORF are listed and annotated in Table S1.

Plasmids were used to transform *Escherichia coli* (*E. coli*) BL21(DE3) wild-type (NEB) or Δ*tig* (John Christodoulou, UCL) cells. Bacteria were grown in ZYM 5052 autoinduction media **(*71*)** with 100 µg/mL ampicillin at 37 °C for 18 hours. Cells were harvested by centrifugation (4,000 rcf, 20 min, 4 °C) and lysed in RNC lysis buffer (Tris-HCl pH 7.5, 150 mM KCl, 10 mM $MgCl_2$) containing 1.25 mg/mL lysozyme, and subjected to 2x freeze-thaw cycles in the presence of 2.72 Kunitz units/µL DNase (Qiagen). The soluble fraction was obtained by centrifugation at 13,200 rpm for 20 min, loaded onto a 35% sucrose cushion in high-salt RNC buffer (50 mM HEPES-KOH pH 7.5, 12 mM $Mg(OAc)_2$, 1 M KOAc, 1 mM dithiothreitol (DTT)), and centrifuged at 4 °C for 2 hours at 250,000 x g (Beckman ultracentrifuge, TLA-110 rotor, tube #363305). The ribosome pellet was resuspended overnight at 4 °C in low-salt RNC buffer (50 mM HEPES-KOH pH 7.5, 12 mM $Mg(OAc)_2$, 100 mM KOAc, 1 mM DTT). The resuspended pellet was applied to in-house prepared GFP-clamp coupled agarose beads (Svend Kjaer, Francis Crick institute) **(*72*)** and incubated overnight at 4 °C. RNCs were eluted by cleavage with 0.4 mg/mL HRV 3C protease (Svend Kjaer, Francis Crick institute) for >5 h, followed by a second round of sucrose cushion ultracentrifugation in high-salt RNC buffer (2 hours, 250,000 x g). For low-salt purifications, both sucrose cushions were prepared using low-salt RNC buffer. In all cases, the final pellet was resuspended in low-salt RNC buffer and A260 was measured to estimate the RNC concentration. See also Fig. S1A.

Where indicated, purified RNCs were incubated with either 50 µg/mL RNaseA (NEB) and 50 mM EDTA, or 2.5 mM puromycin (SantaCruz Biotechnology) at 20 °C for 20 min.

Expression and purification of isolated DHFR variants

The plasmid expressing FL DHFR from a T7 Promoter (supplied as control plasmid with the NEB PURExpress kit) was transformed into *E. coli* BL21(DE3) cells and selected on LB agar plate supplemented with 100 µg/mL ampicillin. A single colony was used to inoculate 500 mL of ZYM 5052 auto induction media containing 100 µg/mL ampicillin. The culture was grown for 24 h in an orbital shaker set to 37 °C. Cells were harvested by centrifugation and resuspended in lysis buffer (25 mM Tris–HCl, pH 7.5, 1 mM ethylenediaminetetraacetic acid (EDTA), and 2 mM DTT). The resuspended cells were supplemented with 1 mM phenylmethanesulfonyl fluoride (PMSF), 0.025 units/mL Benzonase, two tablets of EDTA-free protease inhibitor cocktail, and lysed by sonication. The lysate was centrifuged at 48,000 × g, 4 °C, 45 min, and the supernatant was loaded onto a 6 mL Resource Q column connected to an ÄKTA Pure Protein Purification System. The bound protein was eluted with a linear gradient of 0-500 mM NaCl in lysis buffer. The eluted fractions were examined using NuPAGE 4-12% Bis-Tris gels and the fractions with pure protein were pooled, concentrated using centrifugal filters and injected into a HiLoad 16/600 Superdex 200 pg column that was equilibrated with gel filtration buffer (25 mM Tris–HCl, pH 7.5, 200 mM NaCl, 1 mM EDTA and 1 mM DTT). The fractions with pure protein were pooled, concentrated using centrifugal filters, flash frozen in liquid nitrogen and stored at -80 °C. The concentration of the protein was estimated by absorbance at 280 nm, using an extinction coefficient of 33,585 $M^{-1}.cm^{-1}$.

FL+50$^{stop}$ and 1-126$^{stop}$ were generated from FL+58$^{RNC}$ and 1-126$^{RNC}$, respectively (Table S1) by replacing the SecM sequence with a stop codon using Q5 site-directed mutagenesis (NEB). Proteins were expressed as described above for RNCs. Cells were harvested by centrifugation as above, and the pellet was resuspended in lysis buffer supplemented with 1μL/mL DNase (Roche), 1.25 mg/mL lysozyme ,and 1 protease inhibitor tablet (Roche), then incubated for 30min at 4 °C. Cells were lysed by sonication (3x 1min, 40% amplitude), and the soluble fraction was isolated by centrifugation for 40 min at 20,000 rpm using a JA-25.50 rotor in a Beckman Avanti J-26S XP centrifuge. The supernatant was filtered (0.2 μm) and loaded onto a Resource-Q column connected to ÄKTA Pure Protein Purification System. The protein was eluted using a linear gradient to 50% elution buffer (25mM Tris pH 7.4 at 4 °C, 1mM EDTA, 2 mM DTT, 1 M NaCl) in 20 column volumes. Fractions containing muGFP-DHFR fusion proteins were concentrated and the muGFP-tag cleaved by incubation with 0.4 mg/ml 3C protease overnight at 4 °C. A second, identical round of purification on a RESOURCE-Q column was then performed to separate DHFR and muGFP. Fractions containing cleaved DHFR were pooled, concentrated, and purified from remaining contaminants by size-exclusion chromatography using a Superdex75i column in gel filtration buffer. Fractions containing pure protein were pooled, concentrated using centrifugal filters, flash frozen in liquid nitrogen and stored at -80 °C. Protein concentrations were determined by absorbance at 280 nm

Expression and purification of Trigger factor

Trigger factor (TF) was expressed with a cleavable N-terminal 6xHis-tag from a pPROEX-HTa vector **(*53*)**. TF ΔRBS (F44A/R45A/K46A)**(*47*)** and monomeric TF (V39E, I76E,I80A) **(*55*)** was generated using Q5 site-directed mutagenesis (NEB).

*E. coli* BL21(DE3) cells were transformed with the TF plasmids and selected on LB agar plates supplemented with 100 µg/mL ampicillin. A single colony was used to inoculate 1 L of ZYM 5052 autoinduction media containing 100 µg/mL ampicillin, which was grown overnight at 37 ˚C. The cells were harvested by centrifugation and resuspended in buffer containing 50 mM Tris–HCl pH 7.5, 250 mM NaCl, 10% (v/v) glycerol, 2 mM β-mercaptoethanol, 0.025 units/mL benzonase and 0.1 mM PMSF). The resuspended cells were supplemented with 1 mM PMSF and a tablet of EDTA-free protease inhibitor cocktail and lysed by sonication. The lysate was centrifuged at 48,000 × g for 45 min at 4 °C, and the supernatant was loaded onto a 5 mL HisTrap HP column connected to an ÄKTA Pure Protein Purification System. The bound protein was eluted by washing the column with a linear gradient of 0-500 mM imidazole in lysis buffer. Fractions containing pure protein were pooled, supplemented with 1:100 TEV protease to cleave the His-tag and dialyzed in a buffer containing 20 mM Tris–HCl pH 7.5, 250 mM NaCl, and 2 mM dithiothreitol (DTT) at 4 °C. The protein was passed through a HisTrap column and the flow through containing the cleaved protein was concentrated using centrifugal filters and injected into a Superdex 200i 10/300 GL column that was equilibrated with 20 mM Tris–HCl pH 7.5, 150 mM NaCl, 5% (v/v) glycerol and 1 mM DTT. For monomeric TF, a HiPrep 26/60 Sephacryl S-300 HR column was used for the final SEC step. Fractions containing pure protein were pooled, concentrated using centrifugal filters, flash frozen in liquid nitrogen and stored at -80 °C. The concentration of the protein was estimated using Bradford's assay (***73***).

Mass spectrometry of RNCs

RNCs were purified as described above in either high-salt RNC buffer or, to preserve salt sensitive interactions, in low-salt RNC buffer. Proteins were run for 8 mm using NuPAGE 12% Bis-Tris Gel 1.0 mm 12 wells prior to Coomassie blue staining (Quick Coomassie Stain, Generon). Excised entire 8 mm bands were diced, destained, alkylated and digested with trypsin (modified sequencing grade V5111, Promega). Digests were loaded onto Evotips (Evosep, Odense, Denmark) and tryptic peptides eluted using the "30SPD" gradient via an Evosep One (***74***) HPLC fitted with a 15 cm C18 column (EV1074) into a Lumos Tribrid Orbitrap mass spectrometer (Thermo Scientific) via a nanospray emitter operated at 2200V. The Orbitrap was operated in "Data Dependent Acquisition" mode with precursor ion spectra acquired at 120k resolution in the Orbitrap and MS/MS spectra in the ion trap at 32% HCD collision energy in "TopS" mode. Dynamic exclusion was set to +/- 10 ppm over 15 s, Automatic Gain Control to "standard" and max. injection time to "Dynamic". The vendor's "universal method" was adopted to schedule the ion trap accumulation times. Raw files were processed using Maxquant ((***75***) maxquant.org) and Perseus ((***76***) maxquant.net/perseus) with a recent download of the Uniprot E.Coli reference proteome database together with a common contaminants database. A decoy database of reversed sequences was used to filter false positives, with both peptide and protein false detection rates set to 1%. Quantification of individual *E. coli* proteins was achieved using iBAQ (intensity-based absolute quantification) values normalised to the mean IBAQ value across all ribosomal proteins in each sample. All proteomics data have been deposited to the

ProteomeXchange Consortium via the PRIDE (*77*) partner repository with the dataset identifier PXD036784.

Enzyme activity assays

DHFR activity was measured in low-salt RNC buffer with saturating concentrations of NADPH (Sigma, N6505) and DHF (Sigma, D7006), by following NADPH oxidation via the change in absorbance at 340 nm. For each reaction of 100 μL, 25 nM of DHFR was incubated with 100 μM NADPH for 10 min at 20 °C, then 100 μM DHF was added and the absorbance at 340 nm was immediately recorded for 100 s. To convert the change in absorbance over time into the initial rate, the differential extinction coefficient value of 12.4 $mM^{-1}.cm^{-1}$ was used. All experiments were performed in triplicate at 21°C and measured using a Jasco V-550 Spectrophotometer.

Michaelis-Menten parameters were determined by measuring enzyme activity of 25 nM DHFR as a function of increasing concentration of DHF, keeping NADPH constant at 100 μM. DHF was varied from 0.1 to 30 μM, and the dilutions were made in low-salt RNC buffer for all samples.

Where indicated, 500 nM methotrexate (Sigma A6770) was added to the samples 20 min prior measurement. Where indicated, peptides corresponding to the C-terminus of DHFR were added to the samples and incubated on ice for 40 min prior to adding the substrates. The incubation time was optimized to ensure that the reaction had reached equilibrium. Peptides ($C^{10}$, acetyl-SYCFEILERR-amine or Scr, acetyl-RFIERCELYS-amine) were synthesized by the

Peptide Chemistry Science Technology Platform (Francis Crick Institute) and reconstituted in low-salt RNC buffer.

Pelleting assays

To measure TF binding to RNCs, 10 µM of purified TF was incubated with 1.5-2 µM RNC at 30 °C for 20 min. The reactions were then loaded onto 35% sucrose cushions prepared in either high- or low-salt RNC buffer, and subjected to ultracentrifugation for 2 hours at 250,000 x g (Beckman ultracentrifuge, TLA-100 rotor) to separate unbound TF from the ribosomal fraction. The pellet containing ribosomes was washed once, then resuspended in low-salt RNC buffer and analyzed by SDS-PAGE with Coomassie staining, or immunoblot with antibodies against TF (A01329, GenScript) and small subunit ribosomal protein S2 (abx110548).

Hydrogen/Deuterium exchange mass spectrometry

In addition to the descriptions below, comprehensive experimental details and parameters are provided in Supplemental File Data S1, in the recommended (*78*) tabular format. All HDX MS data have been deposited to the ProteomeXchange Consortium via the PRIDE (*77*) partner repository with the dataset identifier PXD036945. Supplemental File Data S1 contains all the values used to create figures containing HDX MS data.

*Deuterium labeling*

Stock concentrations of purified constructs are listed in Exp Parameters and Replication tab in the Supplemental excel file Data S1. All isolated proteins or RNCs began in storage buffer, as follows. For free FL DHFR and DHFR RNC constructs: 50 mM Hepes-KOH, pH 7.5,

100 mM KOAc, 12 mM $Mg(OAc)_2$, 1 mM DTT; for FL+$50^{stop}$: 25 mM Tris, 1 mM EDTA, 1 mM DTT and 200 mM NaCl; for trigger factor constructs: 20 mM Tris-HCL, pH 7.5, 150 mM NaCl, 5% glycerol, 1 mM DTT; for free ribosomes (obtained from NEB P0763S): 20 mM Hepes-KOH, pH 7.6, 10 mM $Mg(OAc)_2$, 30 mM KCl, 7 mM beta-mercaptoethanol. Proteins were diluted, as needed, to the concentration required for HDX and then labeled with deuterium.

Deuterium labeling was initiated with a 15-fold dilution into labeling buffer (30 µL, 10 mM Hepes-KOH, pD 7.5, 25 mM KOAc, 12 mM $Mg(OAc)_2$, 1 mM DTT, 99.9% $D_2O$). After each labeling time (10 seconds, 100 seconds, and 1000 seconds) at 23 °C, the labeling reaction was quenched with the addition of 20 µL of ice-cold quenching buffer (200 mM potassium phosphate, pH 2.44, 4 M guanidinium chloride, 0.72 M TCEP, $H_2O$), 10 µL 50% immobilized pepsin bead slurry (prepared in house using POROS beads (*79*)) in water with 0.1% formic acid and held on ice for 5 minutes. After on-ice in-solution digestion, the mixture was spun for 15 seconds at 16,000 RCF and 4 °C in Corning® Costar® Spin-X® centrifuge tube filters (Sigma, CLS8163-100EA) then the flow-through was immediately injected into a Waters M-class Acquity UPLC with HDX technology for LC/MS analysis. Undeuterated control samples were prepared for each experiment using the same procedure as outlined above but using 10 mM Hepes-KOH, pH 7.5, 25 mM KOAc, 12 mM $Mg(OAc)_2$, 1 mM DTT, 99.9% $H_2O$ in place of the labeling buffer. Maximally deuterated samples (maxD) were prepared as described in Peterle et al., 2022 (*80*).

*LC-IMS-MS*

The cooling chamber of the UPLC system (based on (*81*), which housed all the chromatographic elements, was held at 0.0 ± 0.1 °C for the entire time of the measurements.

Peptides were trapped and desalted on a VanGuard Pre-Column trap [2.1 mm × 5 mm, ACQUITY UPLC BEH C18, 1.7 µm (Waters, 186002346)] for 3 minutes at 100 µL/min. Peptides were then eluted from the trap using a 5%–35% gradient of acetonitrile over 20 minutes at a flow rate of 100 µL/min, and separated using an ACQUITY UPLC HSS T3, 1.8 µm, 1.0 mm × 50 mm column (Waters, 186003535). The back pressure averaged ~12,950 psi at 0 °C and 5% acetonitrile 95% water, 0.1% formic acid. Mass spectra were acquired using a Waters Synapt G2-Si HDMS$^E$ mass spectrometer in ion mobility (IMS) mode. The mass spectrometer was calibrated with direct infusion of a solution of glu-fibrinopeptide (Sigma, F3261) at 200 femtomole/µL at a flow rate of 5 µL/min prior to data collection. A conventional electrospray source was used, and the instrument was scanned over the range 50 to 2000 m/z. The instrument configuration was the following: capillary was 2.5 kV, trap collision energy at 4 V, sampling cone at 40 V, source temperature of 80 °C and desolvation temperature of 175 °C. All comparison experiments were done under identical experimental conditions such that deuterium levels were not corrected for back-exchange and are therefore reported as relative (*82*). The error of determining the deuterium levels was ± 0.25 Da in this experimental setup.

*HDX MS data processing*

Peptides were identified from replicate HDMS$^E$ analyses (as detailed in the Supplemental File Data S1) of undeuterated control samples using PLGS 3.0.1 (Waters Corporation). Peptide masses were identified from searches using non-specific cleavage of a custom database containing the sequences of each DHFR construct (based on wild-type *E. coli* Uniprot P0ABQ4 that included the linker sequence, see peptide maps in the Supplemental File Data S1), trigger factor (*E. coli* Uniprot P0A850), and all protein sequences in the *E. coli* ribosome as extracted from PDB 4YBB. Searches used the following parameters: no missed cleavages, no PTMs, a low energy threshold of 135, an elevated energy threshold of 35, and an intensity threshold of 500. No false discovery rate (FDR) control was performed. The peptides identified in PLGS (excluding all neutral loss and in-source fragmentation identifications) were then filtered in DynamX 3.0 (Waters Corporation) implementing minimum products per amino acid and consecutive product ions cut-offs described in Exp. Parameters and Replication tab of the Supplemental File Data S1. Peptides meeting the filtering criteria to this point were further processed by DynamX 3.0 (Waters Corporation), including manual inspection of each mass spectrum. The relative amount of deuterium in each peptide was determined with the software by subtracting the centroid mass of the undeuterated form of each peptide from the deuterated form, at each time point, for each condition. These deuterium uptake values were used to generate all uptake graphs and difference maps.

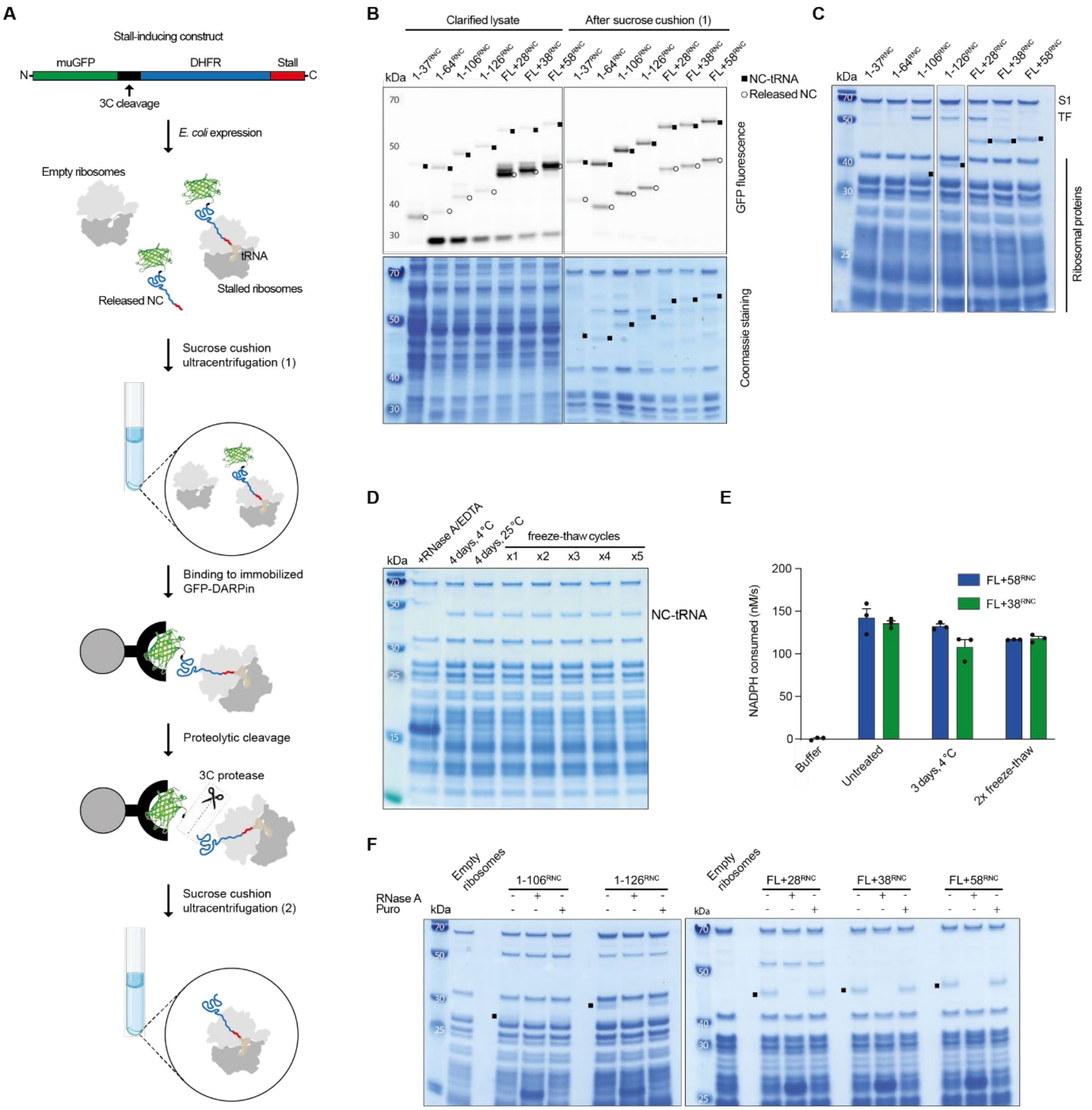


**Fig. S1. Preparation and quality control of stalled ribosome:nascent chain complexes (RNCs).**

**(A)** Overview of RNC purification. Expression constructs include an N-terminal monomeric ultrastable green fluorescent protein (muGFP), followed by a cleavage site for 3C protease, the DHFR sequence of interest, and a C-terminal stalling sequence derived from *M. succiniciproducens* SecM. Expression in *E. coli* results in a mixture of stalled RNCs and free NCs that escape stalling. Total ribosomes are pelleted by sucrose cushion centrifugation, followed by selective purification of RNCs by affinity chromatography using a GFP-binding DARPin with picomolar affinity for GFP. RNCs are then selectively eluted by cleavage with 3C protease, leaving GFP on the resin. A second ultracentrifugation step removes any residual free NC and concentrates the sample.

**(B)** RNCs are tracked in the early stages of the purification by SDS-PAGE followed by either fluorescent imaging of the GFP or Coomassie staining. Ribosome-bound NCs are covalently coupled to peptidyl tRNA (mass ~20 kDa) and can therefore be distinguished from released NCs. Note that shortest RNC ($1\text{-}37^{RNC}$) contains an additional disordered linker after the GFP to improve capture by the DARPin. It therefore migrates more slowly on SDS-PAGE than $1\text{-}64^{RNC}$. Residual released NC, detectable by sensitive fluorescence imaging after the first centrifugation step, is removed by the second round of centrifugation.

**(C)** SDS-PAGE with Coomassie staining of purified RNCs. Ribosomal proteins including S1 are indicated, as is Trigger factor which copurifies with certain RNCs. Where visible between ribosomal proteins, the NC-tRNA band is indicated.

**(D)** RNCs are stable over prolonged incubation and multiple freeze-thaw cycles. The stability of $FL{+}58^{RNC}$ was monitored by the integrity of the NC-tRNA band on SDS-PAGE. Conditions tested included incubation for 4 days at 25 °C or 4 °C, as well as repeated cycles of freezing in liquid $N_2$ followed by thawing on ice. As a positive control, the RNC was destroyed by treatment with 50 μg/mL RNaseA and 50mM EDTA.

**(E)** DHFR in full-length RNCs is stable over prolonged incubation and multiple freeze-thaw cycles. Enzyme activity of FL+38$^{RNC}$ and FL+58$^{RNC}$ was measured either immediately after purification, after incubation at 4 °C for 3 days, or after two freeze-thaw cycles as described in (D).

**(F)** RNCs are resistant to puromycin-induced release. Purified RNCs were treated with either 2.5 mM puromycin or 50 µg/mL RNaseA as a positive control. The NC-tRNA band is indicated in the untreated control lanes.

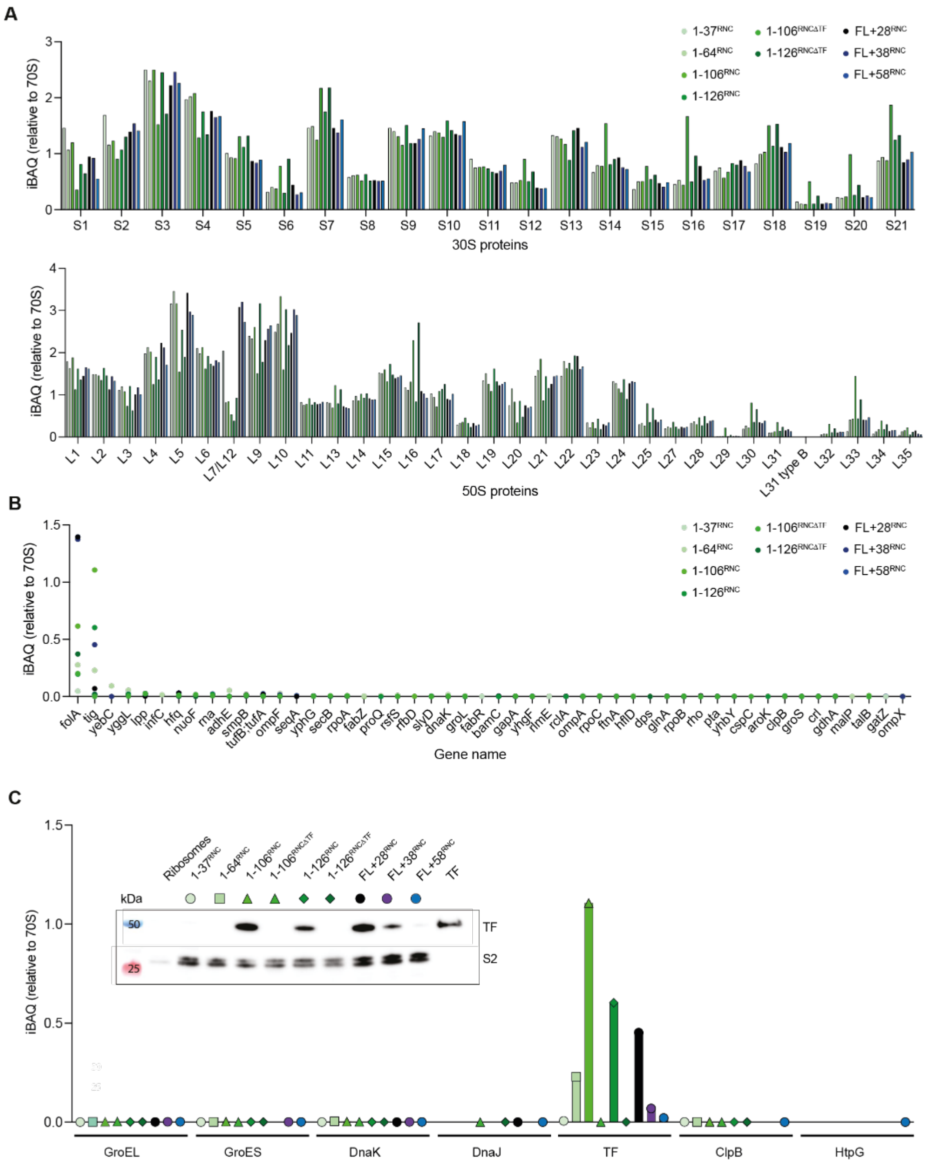


**Fig. S2. Mass spectrometric analysis of RNC composition**

- **(A)** Small (30S) and large (50S) subunit ribosomal proteins detected in purified RNCs. Abundances are based on iBAQ values and normalized to the average iBAQ across all ribosomal proteins in that RNC sample, which is set as 1.

**(B)** Fifty most abundant proteins in RNCs, not including ribosomal proteins. Interactor stoichiometry is calculated as in (A), with the average iBAQ of all detected ribosomal proteins set to 1. iBAQ values for DHFR (*folA*) were not corrected for NC length.

**(C)** Chaperones detected by MS analysis of RNCs. Data are normalized as in (B). The inset shows a western blot of the purified RNCs, with antibodies directed against Trigger factor or 30S protein S2 as a loading control.

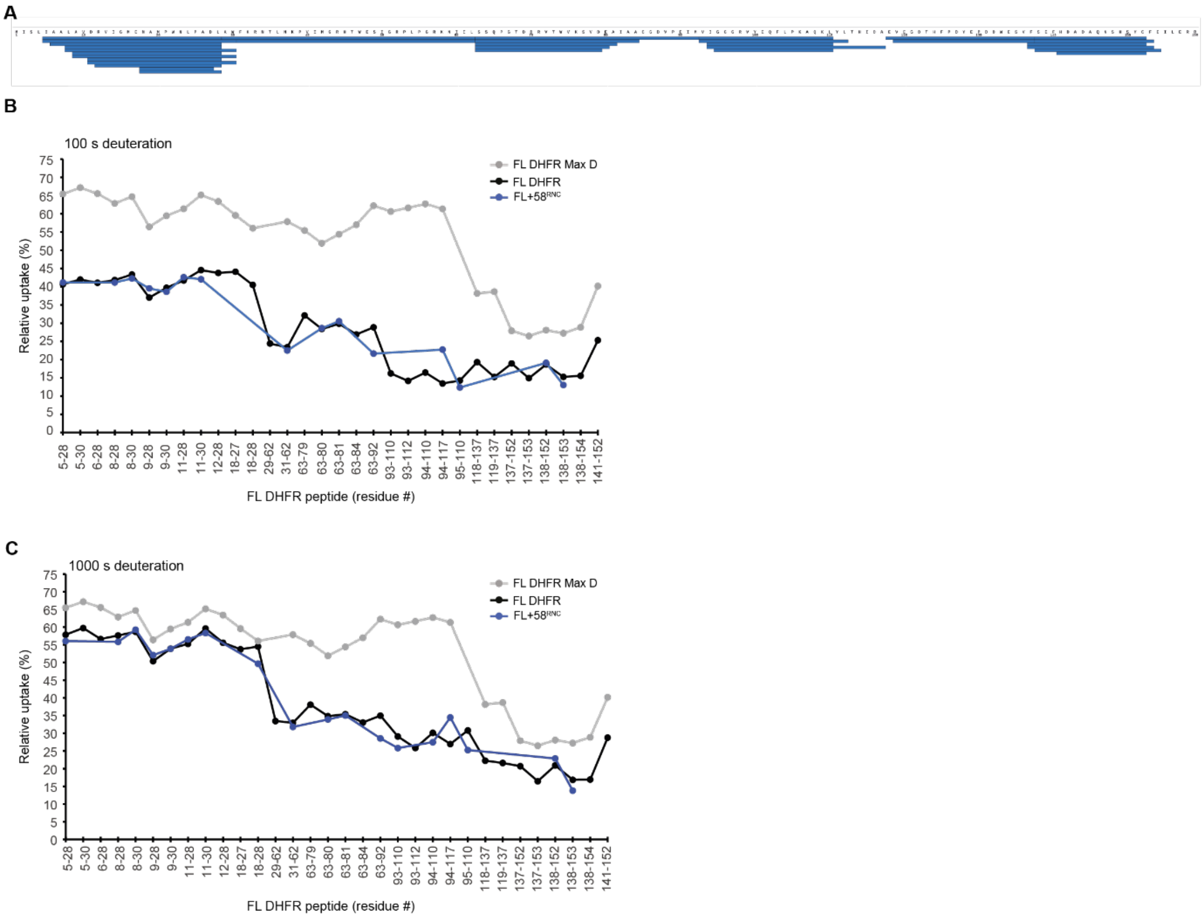


**Fig. S3. HDX MS analysis of full-length DHFR on the ribosome.**

**(A)** Peptide sequence coverage of DHFR in FL+58$^{RNC}$ (34 peptides, 94% coverage). Each peptide is represented by a blue bar.

**(B)** Relative deuterium uptake of DHFR peptides after 100 s exposure to deuterium, as a percentage of the maximum possible exchange, for isolated DHFR (FL DHFR) and FL+58$^{RNC}$. A maximally-deuterated control sample (FL DHFR Max D) is shown as a reference. Values are the average of 2-4 replicates. See also Data S1.

**(C)** As in panel B, but showing relative deuterium uptake of DHFR peptides after 1000 s exposure to deuterium.

Figure S4

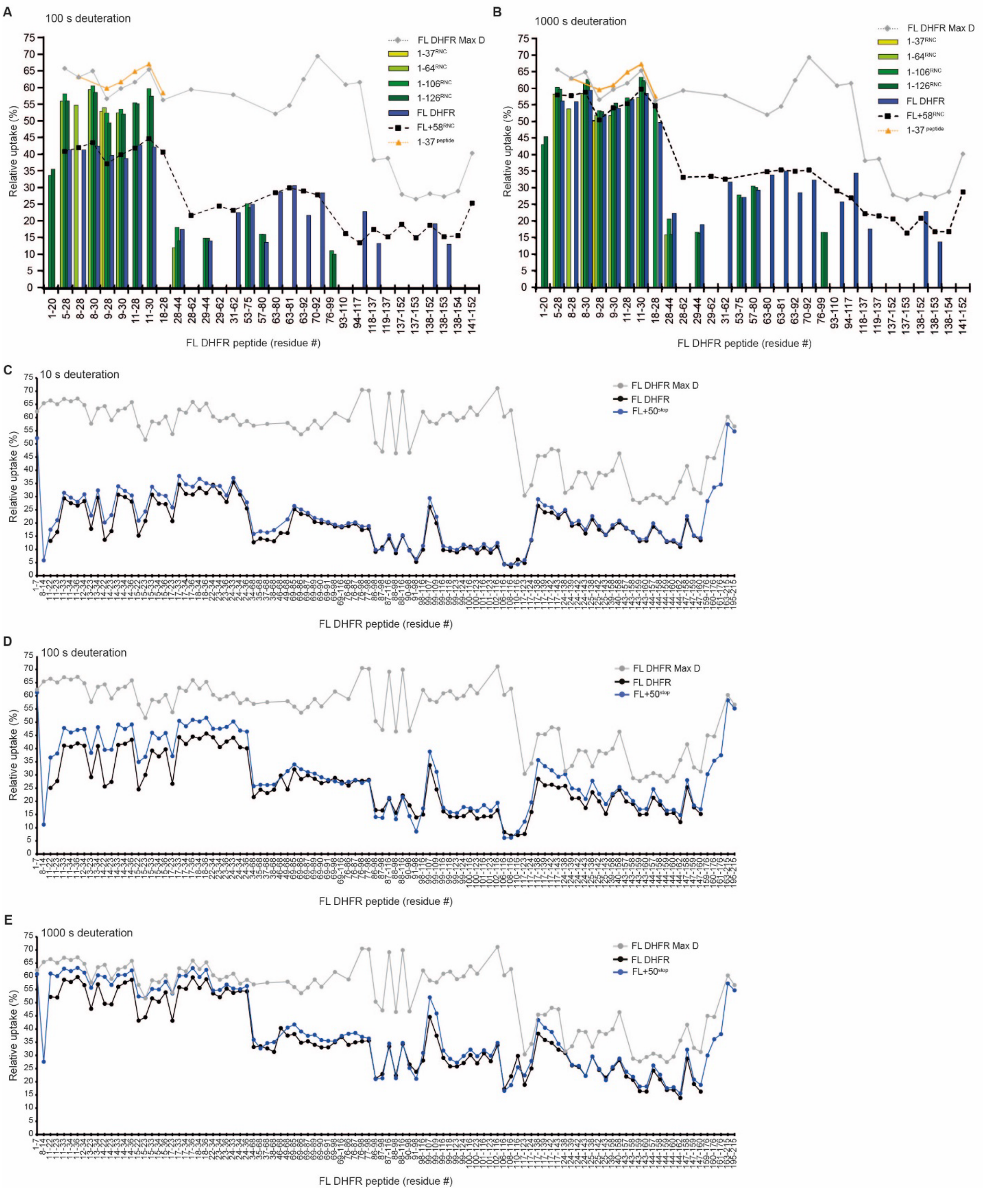


**Fig. S4. HDX MS analysis of DHFR RNCs.**

**(A)** Relative deuterium uptake of DHFR peptides after 100 s exposure to deuterium. Values are the average of 2-4 replicates. See also Data S1.

**(B)** As in panel A, but showing relative deuterium uptake of DHFR peptides after 1000 s exposure to deuterium.

**(C)** Relative deuterium uptake of DHFR peptides after 10 s exposure to deuterium, as a percentage of the maximum possible exchange, for isolated DHFR (FL DHFR) and FL+50$^{stop}$. A maximally-deuterated control sample (FL DHFR Max D) is shown as a reference. Values are the average of 2-4 replicates.

**(D)** As in panel C, but for 100 s exposure to deuterium.

**(E)** As in panel C, but for 1000 s exposure to deuterium. See also Data S1.

Figure S5

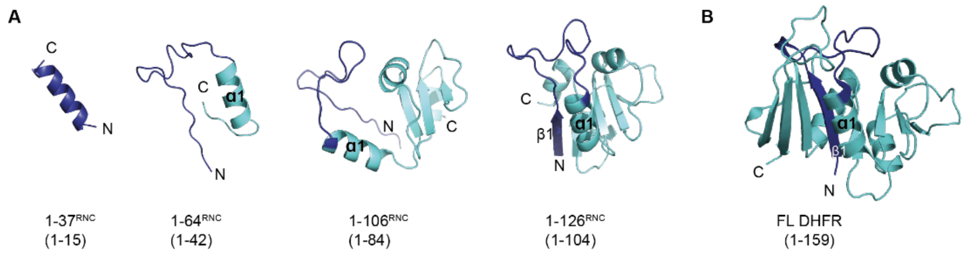


**Fig. S5. Predicted structures of truncated DHFR variants.**

**(A)** Predicted structures of truncated DHFR chains, generated using Alphafold2. Numbers in parenthesis indicate the residues used in structural modelling, which corresponds to the part of each DHFR NC that is expected to be outside the exit tunnel in the RNCs. Residues 1-30, shown to remain unfolded in cotranslational folding intermediates, is colored dark blue. See also Figure 3.

**(B)** Structure of native DHFR (PDB: 5CCC), for comparison to Alphafold2 models. Residues 1-30 are colored dark blue.

Figure S6

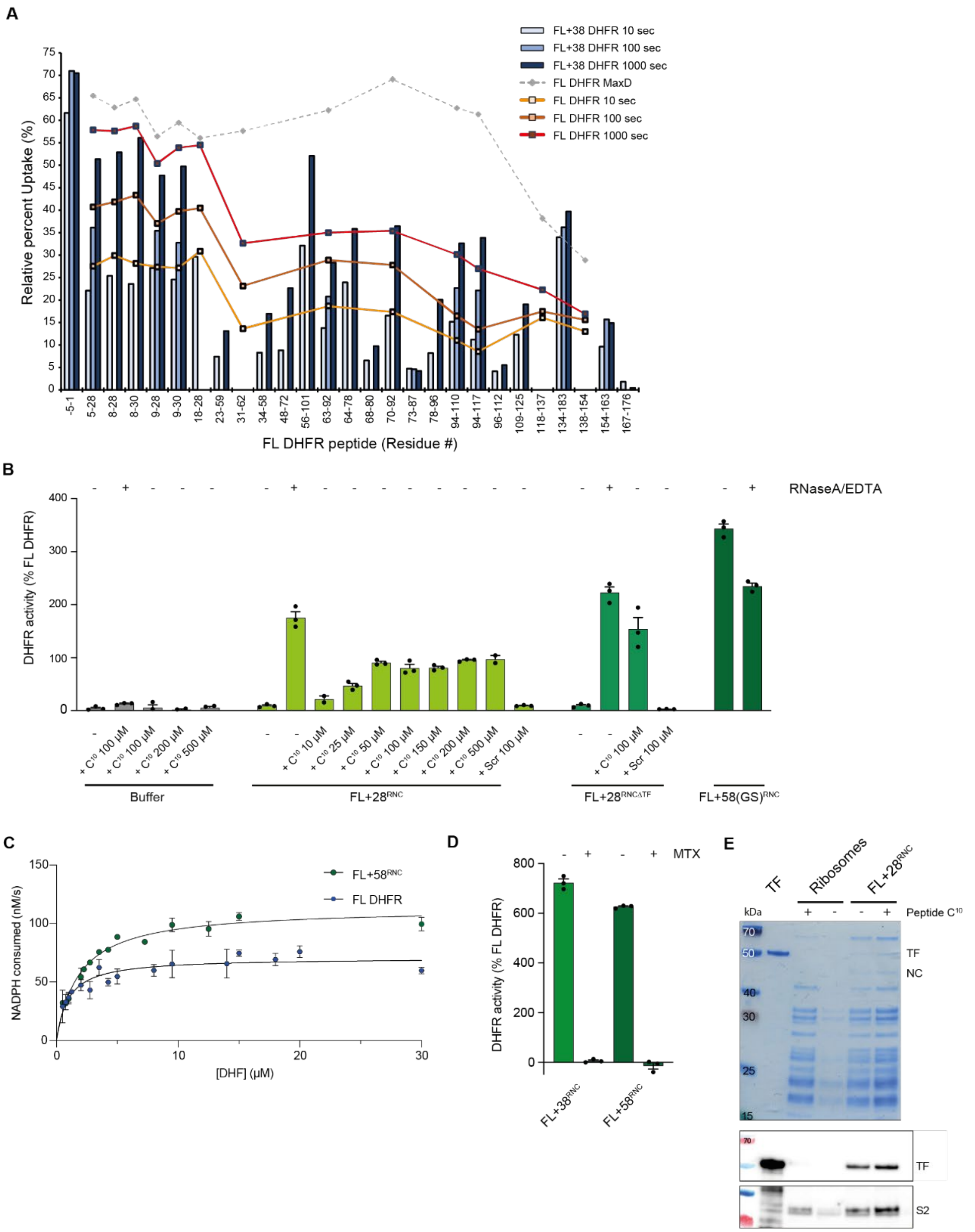


**Fig. S6. Modulation of DHFR activity on the ribosome**

**(A)** Relative deuterium uptake of DHFR peptides after 10 s, 100 s or 1000 s exposure to deuterium, as a percentage of the maximum possible exchange, for isolated DHFR (FL DHFR) and FL+38$^{RNC}$. A maximally-deuterated control sample (FL DHFR Max D) is shown as a reference. Values are the average of 2-4 replicates. See also Data S1.

**(B)** Oxidoreductase activity of 25 nM FL DHFR or RNCs, normalized to the activity of FL DHFR. Where indicated, reactions were supplemented with 50µg/ml RNaseA and 50 mM EDTA, a peptide corresponding to the C-terminus of DHFR ($C^{10}$, SYCFEILERR), or a scrambled-sequence control peptide (Scr, RFIERCELYS). FL+58(GS)$^{RNC}$ is a version of FL+58$^{RNC}$, with the linker between DHFR and the stalling sequence replaced by 25xGS repeats.

**(C)** Michaelis-Menten parameters for DHFR on and off the ribosome. Oxidoreductase activity of 25 nM FL DHFR or FL+58$^{RNC}$, measured at different concentrations of dihydrofolate (DHF). For FL DHFR, $K_M$ = 1.0 µM (95% CI: 0.7 to 1.3); $V_{max}$ = 71 nM/s (95% CI: 66 to 75). For FL+58$^{RNC}$, $K_M$ = 1.9 µM (95% CI: 1.6 to 2.2); $V_{max}$ = 113 nM/s (95% CI: 108 to 118).

**(D)** DHFR RNC activity is sensitive to methotrexate. Oxidoreductase activity of 25 nM FL+38$^{RNC}$ or FL+58$^{RNC}$, with or without 500 nM methotrexate (MTX). Activity is normalized to untreated FL DHFR.

**(E)** Pelleting assay showing TF binding to complemented FL+28$^{RNC}$. Empty ribosomes or FL+28$^{RNC}$ were mixed with 100 µM peptide $C^{10}$ and centrifuged through a high-salt sucrose cushion. The resuspended pellets were analyzed by SDS-PAGE with Coomassie staining, and western blot probing for TF, with S2 was a loading control. Purified TF, not subjected to centrifugation, is shown for reference.

Figure S7

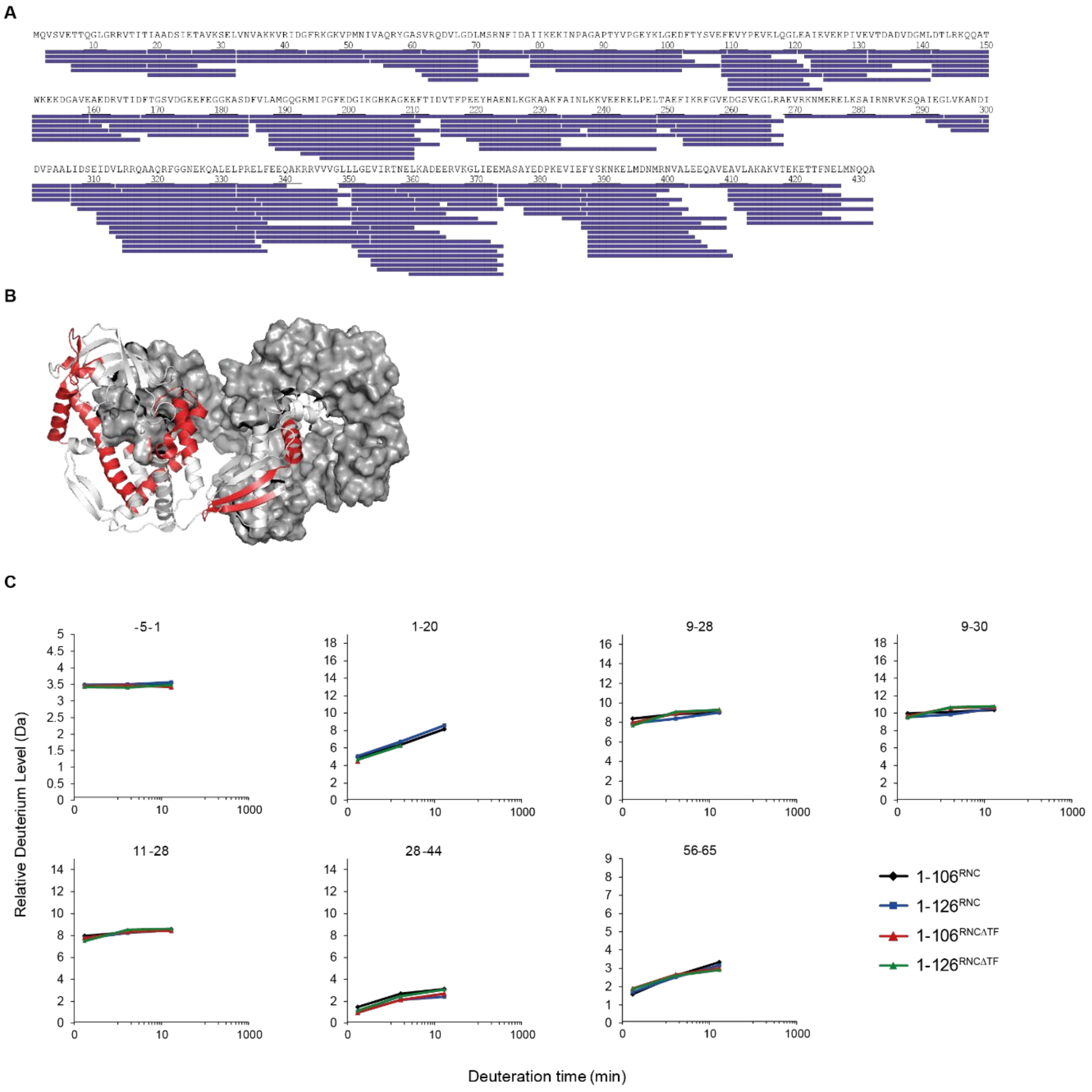


**Fig. S7. HDX MS analysis of TF and ΔTF RNCs.**

**(A)** Peptide sequence coverage of TF in RNCs (181 peptides, 99.5% coverage). Each peptide is represented by a blue bar.

**(B)** Structure of TF dimer (PDB: 6D6S), with peptides that are deprotected in monomeric TF relative to wild-type TF, at any deuteration time point, colored red. One monomer is shown in surface representation. See also Data S1.

**(C)** Relative deuterium uptake as a function of deuteration time for selected peptides in 1-106$^{RNC}$ and 1-126$^{RNC}$, with and without TF. See also Data S1.

Figure S8

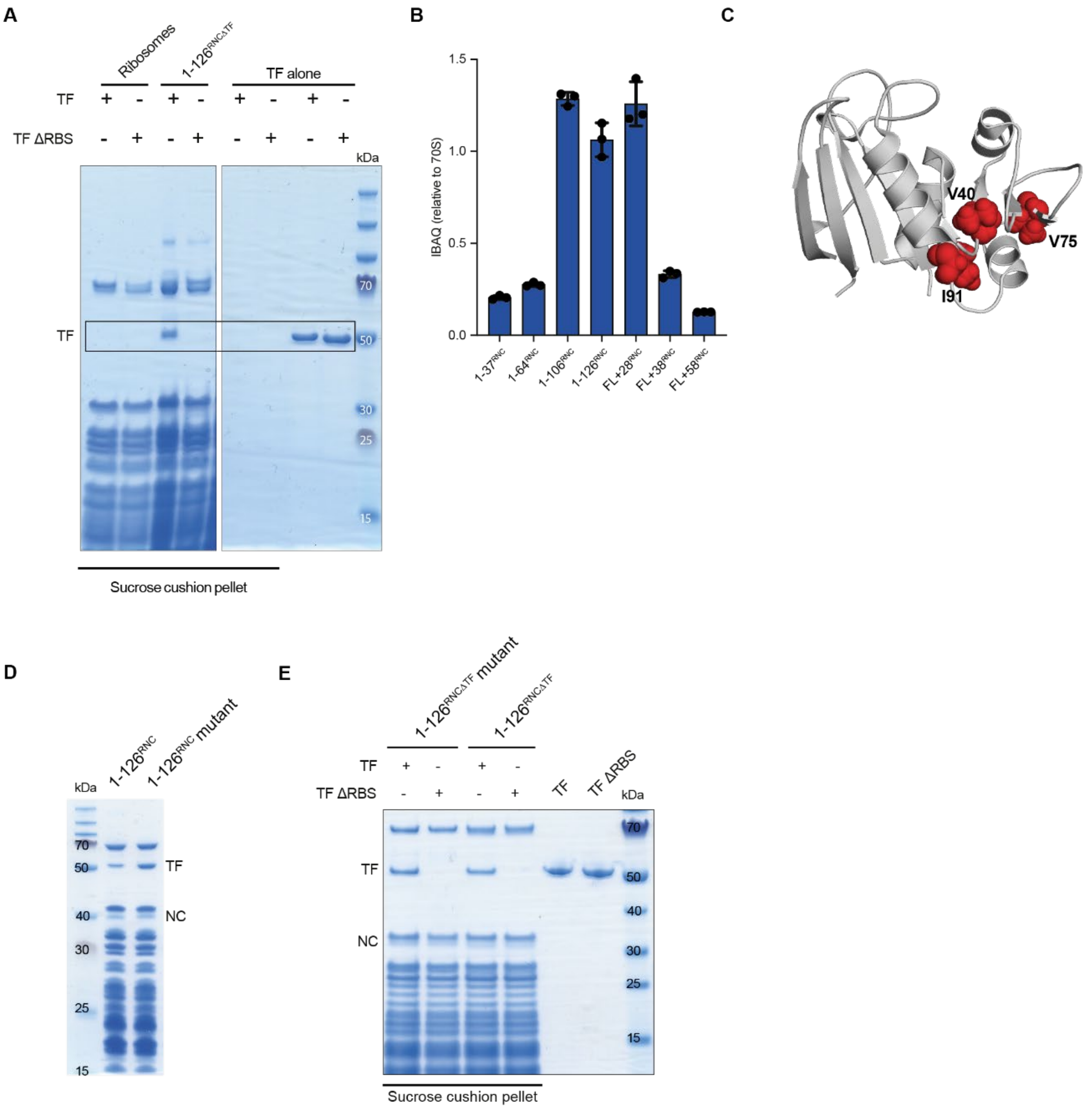


**Fig. S8. Determinants of Trigger factor (TF) binding to RNCs.**

**(A)** Empty 70S ribosomes or 1-126$^{RNC\Delta TF}$ were incubated with wild-type or ribosome-binding-impaired (ΔRBS, F44A/R45A/K46A (*4*)) TF and the reactions were centrifuged through a high-salt (1 M KOAc) sucrose cushion. The pellets were resuspended and analyzed by SDS-PAGE. No TF pelleted in the absence of ribosomes or RNCs.

**(B)** Quantitative proteomic analysis of TF occupancy (as in Fig. S2C) on RNCs purified under low salt (100 mM KOAc) conditions.

**(C)** Structure of DHFR, with the residues mutated in the destabilized variant shown in red.

**(D)** SDS-PAGE of wild-type 1-126$^{RNC}$ and destabilized mutant (1-126$^{RNC}$ mutant) purified under high salt.

**(E)** TF binding to wild-type 1-126$^{RNC\Delta TF}$ and destabilized mutant (1-126$^{RNC\Delta TF}$ mutant) was analyzed as in (A), except that the reactions were centrifuged through a low-salt sucrose cushion.

Figure S9

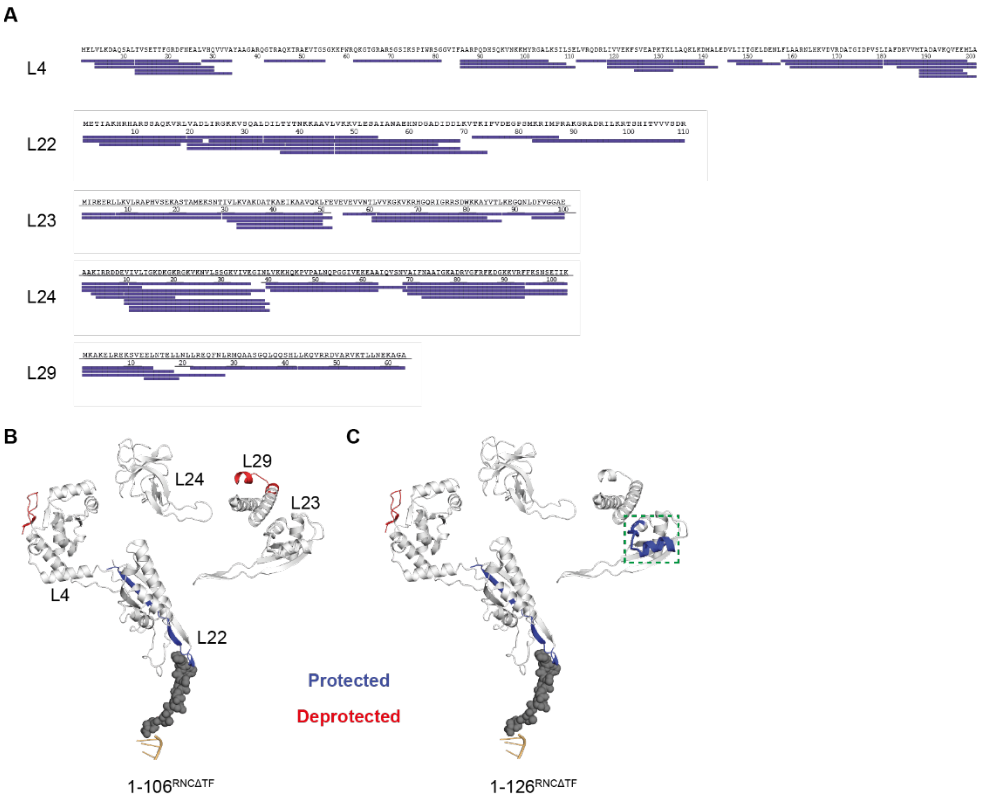


**Fig. S9. HDX MS analysis of ribosomal proteins**

**(A)** Peptide sequence coverage of ribosomal proteins L4 (30 peptides, 90.5% coverage), L22 (17 peptides, 100% coverage), L23 (14 peptides, 98% coverage), L24 (20 peptides, 100% coverage), and L29 (6 peptides, 100% coverage).

**(B)** HDX analysis of ribosomal proteins in 1-106$^{RNC\Delta TF}$. Peptides that are protected from HDX relative to empty ribosomes, at any deuteration time point, are colored blue. Deprotected peptides are colored red.

**(C)** HDX analysis of ribosomal proteins in 1-126$^{RNC\Delta TF}$, as in (B). The TF-docking site on L23 is boxed. See also Data S1.

| Name | DHFR length (aa) | Amino acid sequence | Mw (kDa) |
|---|---|---|---|
| 1-37[RNC] | 1-37 | MSKGEELFTGVVPILVELDGDVNGHKFSVRGEGEGDATNGKLTLKFIC<br>TTGKLPVPWPTLVTTLTYGVLCFSRYPDHMKRHDFFKSAMPEGYVQER<br>TISFKDDGTYKTRAEVKFEGDTLVNRIELKGIDFKEDGNILGHKLEYN<br>FNSHNVYITADKQKNGIKAYFKIRHNVEDGSVQLADHYQQNTPIGDGP<br>VLLPDNHYLSTQSVLSKDPNEKRDHMVLLEDVTAAGITHGMDELYK**GS**<br>**GS**LEVLFQGP**GSGSGSGSGSGSGSGSGSGSGSGSGSENLYFQGSG**MIS<br>LIAALAVDRVIGMENAMPWNLPADLAWFKRNTLNWWWPRIRGPPGS | 8.5 |
| 1-64[RNC] | 1-64 | MSKGEELFTGVVPILVELDGDVNGHKFSVRGEGEGDATNGKLTLKFIC<br>TTGKLPVPWPTLVTTLTYGVLCFSRYPDHMKRHDFFKSAMPEGYVQER<br>TISFKDDGTYKTRAEVKFEGDTLVNRIELKGIDFKEDGNILGHKLEYN<br>FNSHNVYITADKQKNGIKAYFKIRHNVEDGSVQLADHYQQNTPIGDGP<br>VLLPDNHYLSTQSVLSKDPNEKRDHMVLLEDVTAAGITHGMDELYK**GS**<br>**GS**LEVLFQGP**GSGS**MISLIAALAVDRVIGMENAMPWNLPADLAWFKRN<br>TLNKPVIMGRHTWESIGRPLPGRKNIILSSWWWPRIRGPPGS | 9 |
| 1-106[RNC] | 1-106 | MSKGEELFTGVVPILVELDGDVNGHKFSVRGEGEGDATNGKLTLKFIC<br>TTGKLPVPWPTLVTTLTYGVLCFSRYPDHMKRHDFFKSAMPEGYVQER<br>TISFKDDGTYKTRAEVKFEGDTLVNRIELKGIDFKEDGNILGHKLEYN<br>FNSHNVYITADKQKNGIKAYFKIRHNVEDGSVQLADHYQQNTPIGDGP<br>VLLPDNHYLSTQSVLSKDPNEKRDHMVLLEDVTAAGITHGMDELYK**GS**<br>**GS**LEVLFQGP**GSGS**MISLIAALAVDRVIGMENAMPWNLPADLAWFKRN<br>TLNKPVIMGRHTWESIGRPLPGRKNIILSSQPGTDDRVTWVKSVDEAI<br>AACGDVPEIMVIGGGRVYEQFLPKWWWPRIRGPPGS | 13.5 |
| 1-126[RNC] | 1-126 | MSKGEELFTGVVPILVELDGDVNGHKFSVRGEGEGDATNGKLTLKFIC<br>TTGKLPVPWPTLVTTLTYGVLCFSRYPDHMKRHDFFKSAMPEGYVQER<br>TISFKDDGTYKTRAEVKFEGDTLVNRIELKGIDFKEDGNILGHKLEYN<br>FNSHNVYITADKQKNGIKAYFKIRHNVEDGSVQLADHYQQNTPIGDGP<br>VLLPDNHYLSTQSVLSKDPNEKRDHMVLLEDVTAAGITHGMDELYK**GS**<br>**GS**LEVLFQGP**GSGS**MISLIAALAVDRVIGMENAMPWNLPADLAWFKRN<br>TLNKPVIMGRHTWESIGRPLPGRKNIILSSQPGTDDRVTWVKSVDEAI<br>AACGDVPEIMVIGGGRVYEQFLPKAQKLYLTHIDAEVEGDTHFPWWWP<br>RIRGPPGS | 15.5 |
| 1-126[RNC] mutant (V40A, V75H, I91L) | 1-126 | MSKGEELFTGVVPILVELDGDVNGHKFSVRGEGEGDATNGKLTLKFIC<br>TTGKLPVPWPTLVTTLTYGVLCFSRYPDHMKRHDFFKSAMPEGYVQER<br>TISFKDDGTYKTRAEVKFEGDTLVNRIELKGIDFKEDGNILGHKLEYN<br>FNSHNVYITADKQKNGIKAYFKIRHNVEDGSVQLADHYQQNTPIGDGP<br>VLLPDNHYLSTQSVLSKDPNEKRDHMVLLEDVTAAGITHGMDELYK**GS**<br>**GS**LEVLFQGP**GSGS**MISLIAALAVDRVIGMENAMPWNLPADLAWFKRN<br>TLNKPAIMGRHTWESIGRPLPGRKNIILSSQPGTDDRVTWHKSVDEAI<br>AACGDVPELMVIGGGRVYEQFLPKAQKLYLTHIDAEVEGDTHFPWWWP<br>RIRGPPGS | 15.5 |
| FL+28[RNC] | 1-159 | MSKGEELFTGVVPILVELDGDVNGHKFSVRGEGEGDATNGKLTLKFIC<br>TTGKLPVPWPTLVTTLTYGVLCFSRYPDHMKRHDFFKSAMPEGYVQER<br>TISFKDDGTYKTRAEVKFEGDTLVNRIELKGIDFKEDGNILGHKLEYN<br>FNSHNVYITADKQKNGIKAYFKIRHNVEDGSVQLADHYQQNTPIGDGP<br>VLLPDNHYLSTQSVLSKDPNEKRDHMVLLEDVTAAGITHGMDELYK**GS**<br>**GS**LEVLFQGP**GSGS**MISLIAALAVDRVIGMENAMPWNLPADLAWFKRN<br>TLNKPVIMGRHTWESIGRPLPGRKNIILSSQPGTDDRVTWVKSVDEAI<br>AACGDVPEIMVIGGGRVYEQFLPKAQKLYLTHIDAEVEGDTHFPDYEP<br>DDWESVFSEFHDADAQNSHSYCFEILERR**GSGGDLGSGGYLGSGGLDG**<br>**SGGVEGSGGGFL**WWWPRIRGPPGS | 21.4 |
| FL+38[RNC] | 1-159 | MSKGEELFTGVVPILVELDGDVNGHKFSVRGEGEGDATNGKLTLKFIC<br>TTGKLPVPWPTLVTTLTYGVLCFSRYPDHMKRHDFFKSAMPEGYVQER<br>TISFKDDGTYKTRAEVKFEGDTLVNRIELKGIDFKEDGNILGHKLEYN | 22.3 |

| | | | |
|---|---|---|---|
| | | FNSHNVYITADKQKNGIKAYFKIRHNVEDGSVQLADHYQQNTPIGDGP<br>VLLPDNHYLSTQSVLSKDPNEKRDHMVLLEDVTAAGITHGMDELYK**GS**<br>**GS**LEVLFQGP**GSGS**MISLIAALAVDRVIGMENAMPWNLPADLAWFKRN<br>TLNKPVIMGRHTWESIGRPLPGRKNIILSSQPGTDDRVTWVKSVDEAI<br>AACGDVPEIMVIGGGRVYEQFLPKAQKLYLTHIDAEVEGDTHFPDYEP<br>DDWESVFSEFHDADAQNSHSYCFEILERRGSGGDLGSGGYLGSGGLDG<br>SGGVEGSGGFLWWWPRIRGPPGS | |
| FL+58$^{RNC}$ | 1-159 | MSKGEELFTGVVPILVELDGDVNGHKFSVRGEGEGDATNGKLTLKFIC<br>TTGKLPVPWPTLVTTLTYGVLCFSRYPDHMKRHDFFKSAMPEGYVQER<br>TISFKDDGTYKTRAEVKFEGDTLVNRIELKGIDFKEDGNILGHKLEYN<br>FNSHNVYITADKQKNGIKAYFKIRHNVEDGSVQLADHYQQNTPIGDGP<br>VLLPDNHYLSTQSVLSKDPNEKRDHMVLLEDVTAAGITHGMDELYK**GS**<br>**GS**LEVLFQGP**GSGS**MISLIAALAVDRVIGMENAMPWNLPADLAWFKRN<br>TLNKPVIMGRHTWESIGRPLPGRKNIILSSQPGTDDRVTWVKSVDEAI<br>AACGDVPEIMVIGGGRVYEQFLPKAQKLYLTHIDAEVEGDTHFPDYEP<br>DDWESVFSEFHDADAQNSHSYCFEILERR**GSGGDLGSGGYLGSGGLDG**<br>**SGGVEGSGGFLGSGGLDGSGGLYGSGGVEGS**WWWPRIRGPPGS | 23.9 |
| FL+58(GS)$^{RNC}$ | 1-159 | MSKGEELFTGVVPILVELDGDVNGHKFSVRGEGEGDATNGKLTLKFIC<br>TTGKLPVPWPTLVTTLTYGVLCFSRYPDHMKRHDFFKSAMPEGYVQER<br>TISFKDDGTYKTRAEVKFEGDTLVNRIELKGIDFKEDGNILGHKLEYN<br>FNSHNVYITADKQKNGIKAYFKIRHNVEDGSVQLADHYQQNTPIGDGP<br>VLLPDNHYLSTQSVLSKDPNEKRDHMVLLEDVTAAGITHGMDELYK**GS**<br>**GS**LEVLFQGP**GSGS**MISLIAALAVDRVIGMENAMPWNLPADLAWFKRN<br>TLNKPVIMGRHTWESIGRPLPGRKNIILSSQPGTDDRVTWVKSVDEAI<br>AACGDVPEIMVIGGGRVYEQFLPKAQKLYLTHIDAEVEGDTHFPDYEP<br>DDWESVFSEFHDADAQNSHSYCFEILERR**GSGSGSGSGSGSGSGSGSG**<br>**SGSGSGSGSGSGSGSGSGSGSGSGSGSGSGS**WWWPRIRGPPGS | 23.9 |

**Table S1. DHFR RNC sequences**. The N-terminal muGFP (green), functioning as an affinity tag, precedes a 3C cleavage sequence (blue) that is flanked on either side by $(GS)_2$ linkers (bold). The subsequent DHFR sequence fragment is followed by a C-terminal stall-inducing sequence derived from *M. succiniciproducens* SecM (33). Structural analysis of ribosomes stalled on wild-type *E. coli* SecM indicates that various stalling scenarios are possible (59). When describing our RNCs we assume that the ribosome stalls at the first available opportunity, with the penultimate $tRNA^{Pro}$ in the A-site and the first 8 residues from the stall-inducing sequence occupying the exit tunnel. In the full-length RNCs an additional disordered linker precedes the stall sequence (bold). In 1-37$^{RNC}$, the GS linker following the 3C cleavage sequence is extended to improve capture by the GFP affinity resin. Sites of point mutations are underlined.

**Data S1. (separate file)**

**Hydrogen-deuterium exchange-mass spectrometry (HDX MS) data.**

Consisting of the following tabs in Microsoft Excel:

- Experimental parameters and replication
- Data in Figures 2B; S3
- Data in Figures 3A,B; S4A,B
- Data in Figure 3D,E; S4C-E
- Data in Figures 4A; S6A
- Data in Figures 5B; S7A,B
- Data in Figure S7C
- Data in Figures 6; S9
